# A spatial and cellular distribution of neurotropic virus infection in the mouse brain revealed by fMOST and single cell RNA-seq

**DOI:** 10.1101/2021.03.26.436691

**Authors:** Yachun Zhang, Xudong Xing, Ben Long, Yandi Cao, Simeng Hu, Xiangning Li, Yalan Yu, Dayong Tian, Baokun Sui, Zhaochen Luo, Wei Liu, Lei Lv, Qiong Wu, Jinxia Dai, Ming Zhou, Heyou Han, Zhen F. Fu, Hui Gong, Fan Bai, Ling Zhao

## Abstract

Neurotropic virus infection can cause serious damage to the central nervous system (CNS) in both human and animals. The complexity of the CNS poses unique challenges to investigate the infection of these viruses in the brain using traditional techniques. In this study, we explore the use of fluorescence micro-optical sectioning tomography (fMOST) and single cell RNA sequencing (scRNA-seq) to map the spatial and cellular distribution of a representative neurotropic virus, rabies virus (RABV), in the whole brain. Mice were inoculated with a lethal dose of recombinant RABV expressing enhanced green fluorescent protein (EGFP) under different infection routes, and a three-dimensional view of the distribution of RABV in the whole mouse brain was obtained using fMOST. Meanwhile, we pinpointed the cellular distribution of RABV by utilizing scRNA-seq. Our fMOST data provide the first evidence that RABV can infect multiple nuclei related to fear independent of different infection routes. More surprisingly, scRNA-seq data indicate that besides neurons RABV can infect macrophages and NK cells *in vivo*. Collectively, this study draws a comprehensively spatial and cellular map of RABV infection in the mouse brain, providing a novel and insightful strategy to investigate the pathogenesis of neurotropic viruses.

## 1. Introduction

Neurotropic virus infection can cause an array of immediate and delayed neuropathology in human and animals, and nearly half of the emerging viruses can invade the CNS ^1^. These infections pose a major challenge to human and animal healthcare worldwide. The complex structures and functions of the CNS and the diversity of neurotropic viruses make it difficult to find an effective treatment of these diseases. By combining human pathological data with experimental animal models, virologists have much advanced our understanding of the mechanisms underlying how viruses enter the CNS and cause neurological disease, but more in-depth studies are still in urgent need to facilitate the development of novel vaccines and antiviral therapeutics ^2^. Rabies virus (RABV) is a typical neurotropic virus belonging to *Rhabdoviridae* family. RABV hijacks the cellular transport machinery and moves along microtubules by retrograde axonal transport to the nearest sensory neurons in the dorsal root ganglion or anterior horn of the spinal cord ^3^. After replication, RABV travels along the corticospinal tract to the brain where the virus can efficiently replicate and infect in most regions, resulting in fatal encephalitis ^4^. However, due to the complexity of the CNS, the exact regions and cell types infected by RABV are still kept elusive, impeding the endeavor of researchers to find the therapeutic target for RABV.

For many decades, light microscopy was the key tool for investigating the invasion and spread of neurotropic viruses in the host’s brain. However, to observe a whole animal brain, which is centimeter-sized, is beyond the view field of modern light microscopy techniques including confocal and multiphoton microscopy. Histological sections were usually prepared for observing the internal microstructures of large specimens, but it is difficult to align the serial figures of continuous ultrathin sectioning on large specimens by a traditional light microscopy. In 2010, a micro-optical sectioning tomography (MOST) system was developed, which allowed the mapping of a whole mouse brain to the single neuron level^5^. In 2013, a fluorescence MOST (fMOST) was developed by using a resin-embedding method for maintaining fluorescence and an automated fluorescence MOST system for long-term stable imaging ^6^. Recently, a modified fMOST, benefiting from simple sample preparation and high-throughput imaging, was applied to determine the detailed molecular characteristics of neural circuits ^7, 8^. This fMOST system provides wide-field optical-sectioning imaging of an agarose-embedded sample with high efficiency. Single-cell sequencing is an emerging technology that allows high-throughput sequencing analyses of genome, transcriptome, and methylome at the single-cell level. It has been recently implemented in the study of influenza virus ^9^, HIV ^10^, Ebola virus^11^ and SARS-CoV-2 ^12^ infections, providing an unbiased and comprehensive visualization of the target cells that virus can infect and a global immunological responses of the host.

In this study, we constructed a recombinant RABV expressing EGFP (RABV-EGFP) and utilized the fMOST system to portray the spatial distribution of RABV in the brain of infected mice. Meanwhile, scRNA-seq was applied to pinpoint the cellular distribution of RABV infection. These two combined techniques reveal a three-dimension (3D) distribution of RABV in the whole brain and identify several major cell types infected by RABV. Our results provide a better understanding of the pathogenesis of neurotropic viruses and shed new light on the development of novel therapies for their infection.

## 2. Results

### 2.1. Brain-wide spatial mapping of RABV infection

To map the spatial distribution of RABV in the whole brain, we constructed and characterized a recombinant RABV expressing EGFP (RABV-EGFP) *in vitro* and *in vivo* (Figure S1). Then we infected C57BL/6 mice with RABV-EGFP and investigated its spread in the brain by using fMOST techniques. The research strategy was briefly depicted in Figure 1A. To evaluate the impact of infection routes on the spatial distribution of RABV in brains, groups of C57BL/6 mice were infected with different dilutions of RABV-EGFP via three different routes, including intramuscular (i.m.) injection in the hind limbs, subcutaneous (o.s.) inoculation under the ears and intranasal administration (i.n.), and the median lethal dose (LD_50_) of RABV-EGFP for each infection method was calculated (Figure S2). Then the mice were infected with 10×LD_50_ of RABV-EGFP via three different routes. At the moribund stage, the mice were euthanized and the brains were harvested. Under each infection route, the whole brain was extensively infected by RABV and the viral load was almost comparable in cerebrum, cerebellum and brain stem analyzed by quantitative real-time PCR (qPCR) (Figure S3).

**Figure.1.**
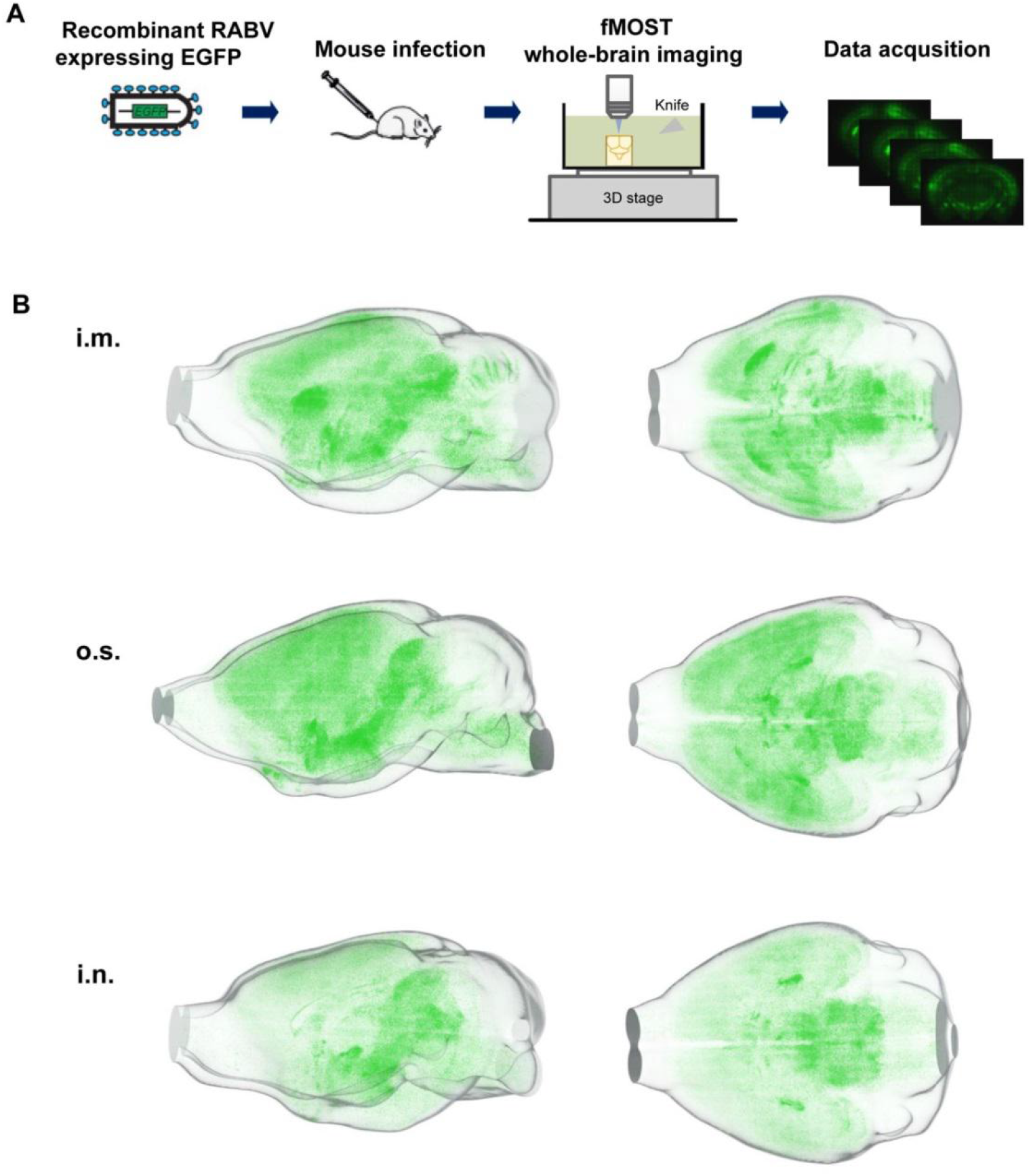
The three-dimension (3D) distribution of RABV in the whole mouse brain. (A) Scheme of the experimental workflow for mapping the 3D distribution of RABV in the whole mouse brain using the fMOST technique. (B) Groups of C57BL/6 mice were inoculated with 10×LD_50_ of the recombinant RABV expressing EGFP, RABV-EGFP, by the intramuscular (i.m.), the otic subcutaneous (o.s.), or the intranasal (i.n.) route. At the moribund stage, the brains were harvested and prepared for fMOST processing. Representative pictures of the distribution of RABV in a whole mouse brain by vertical and sagittal views are shown. The overall distribution of RABV in the mouse brain can be viewed in supplementary video 1-3.

For fMOST analysis, the oxidized agarose-embedded brains were prepared for full-volumetric imaging with a resolution of 0.32 μm × 0.32 μm × 5 μm by automatically repeating sectioning-imaging cycles. Approximately 3,000 equidistant coronal sections composed a whole-brain dataset, and the self-registration of the dataset allowed for convenient 3D visualization of the distribution of RABV-EGFP. Corresponding figures from the reference channel were used to assist the identification of brain contours and areas. To ensure the reproducibility of the technique and the veracity of the analysis, three mouse brains from each infection route were imaged. As shown in Figure 1B, we found that RABV was predominant in the cortex, hypothalamus, hippocampus, midbrain, and brain stem. Specially, mice inoculated by i.m. infection had virus concentrated in the motor area of the cerebral cortex and the head of striatum, while mice inoculated by i.n. infection had virus primarily in dentate gyrus. The extensive RABV infection along cerebellar gyri could be observed only post i.m. inoculation. Together, these findings suggest that the spatial distribution of RABV in the brain is influenced by the route of infection.

### 2.2. The brain regions infected by RABV

To further identify the brain regions infected by RABV under different inoculation routes, images of representative coronal sections, spanning from olfactory bulb to medulla, were selected and examined. The anatomical localization of RABV in these sections is shown in Figure 2A and the abbreviations of anatomical structures can be found in Table 1. Based on cellular morphology, we found that most of RABV-infected cells were neurons, which is consistent with previous findings ^4, 13^. To consolidate this finding, we stained RABV P protein (RABV-P) and neuron marker, NeuN, in the mouse brain sections, and observed them under an immunofluorescence microscope. The results showed strong co-localization between RABV-P and neurons (Figure S4). To illustrate infections in a single cell, enlarged views of the motor cortex (MO), the bed nuclei of the stria terminalis (BNST), and the superior colliculus (SC) are shown in Figure 2B.

**Figure.2.**
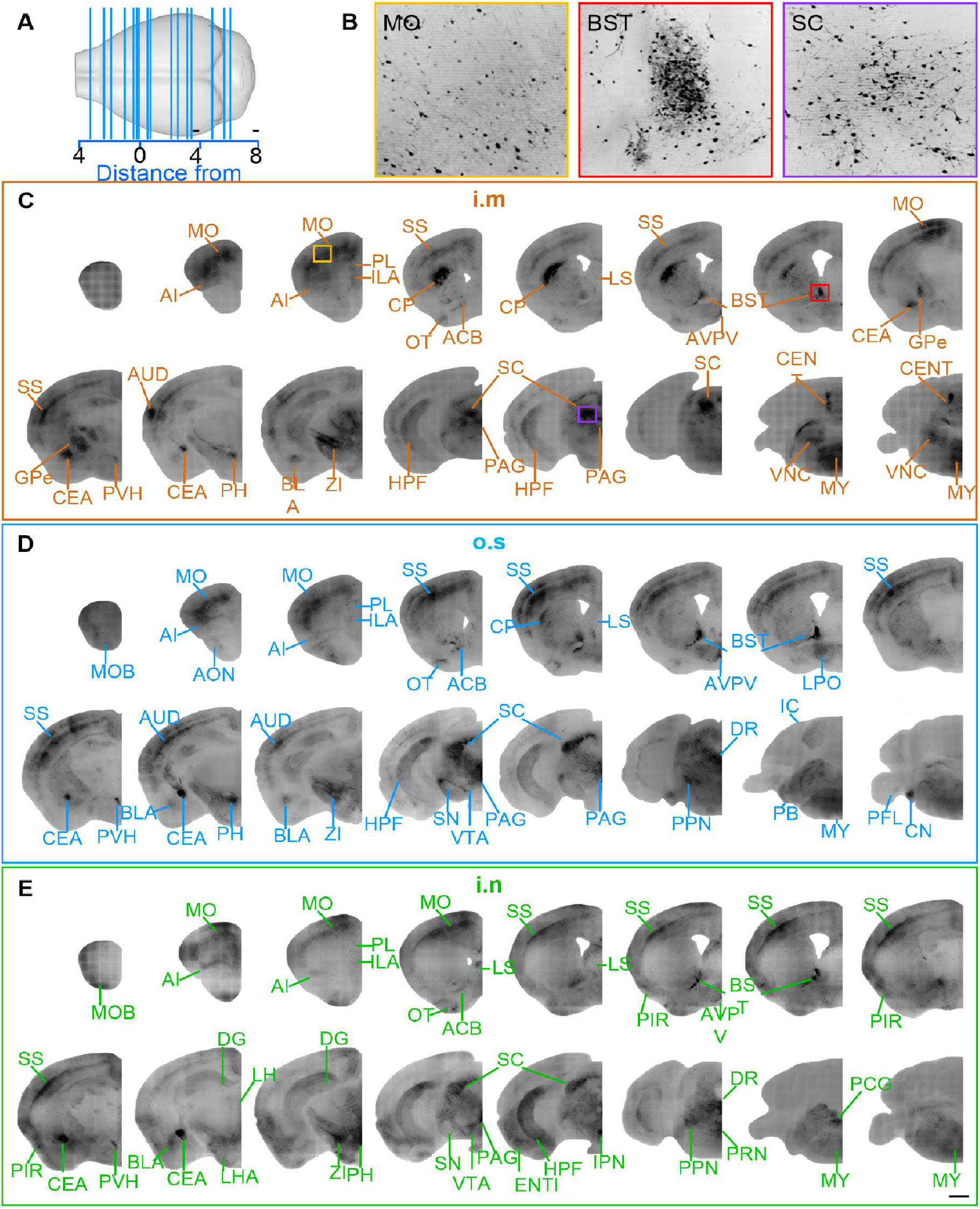
Anatomical classification of RABV distribution by different infection routes. (A) The anatomical localization of the selected coronal sections shown in (C-E). Scale bar, 1 mm. (B) Enlarged views, indicated by a yellow, red, and purple box, respectively, of the motor cortex (MO), bed nuclei of the stria terminalis (BNST), and the superior colliculus (SC) from panel C. A single RABV-infected cell is shown. Scale bar, 100 μm. (C-E) The distribution of RABV by infection route, i.m., o.s., and i.n., respectively.

**Table 1.**
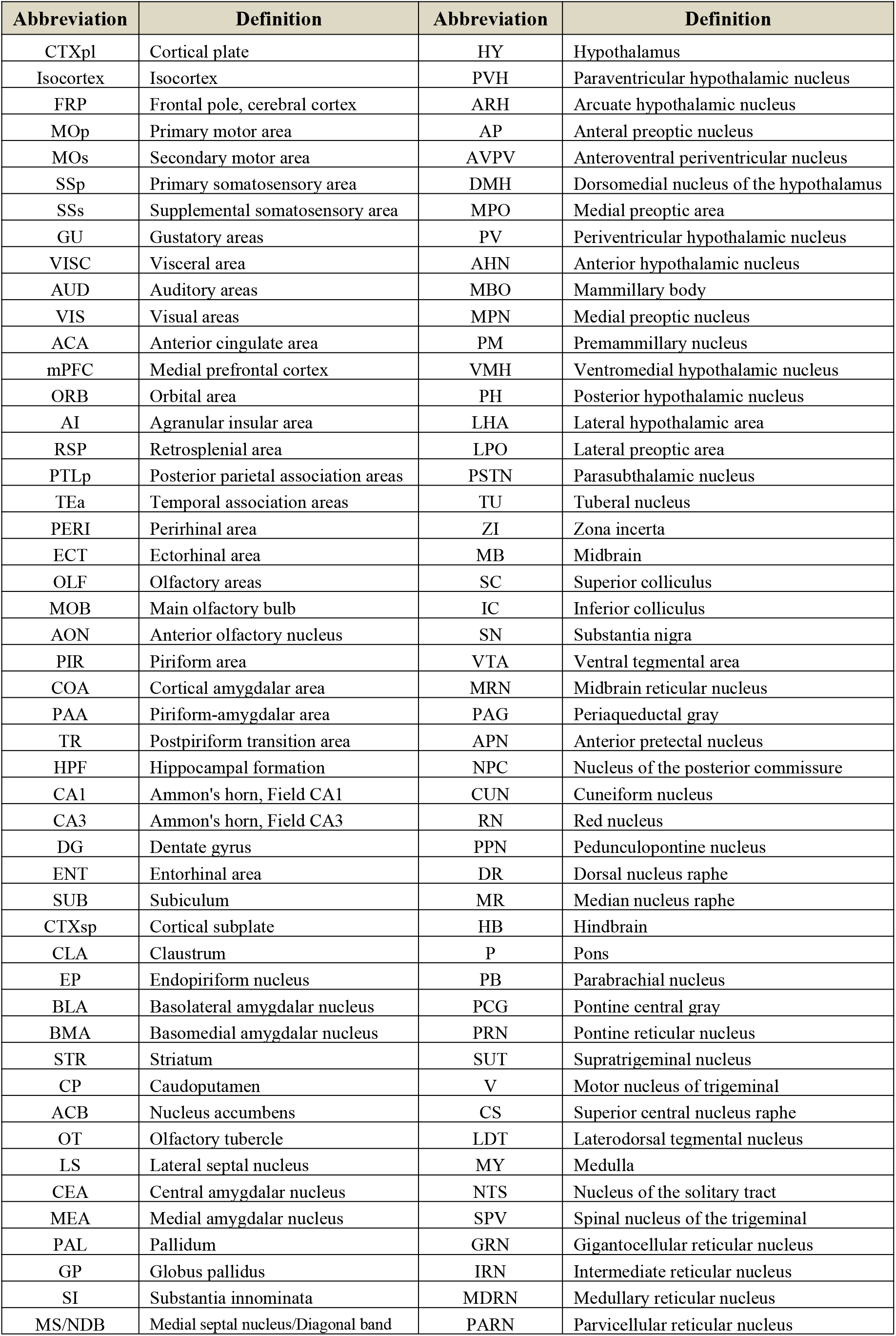
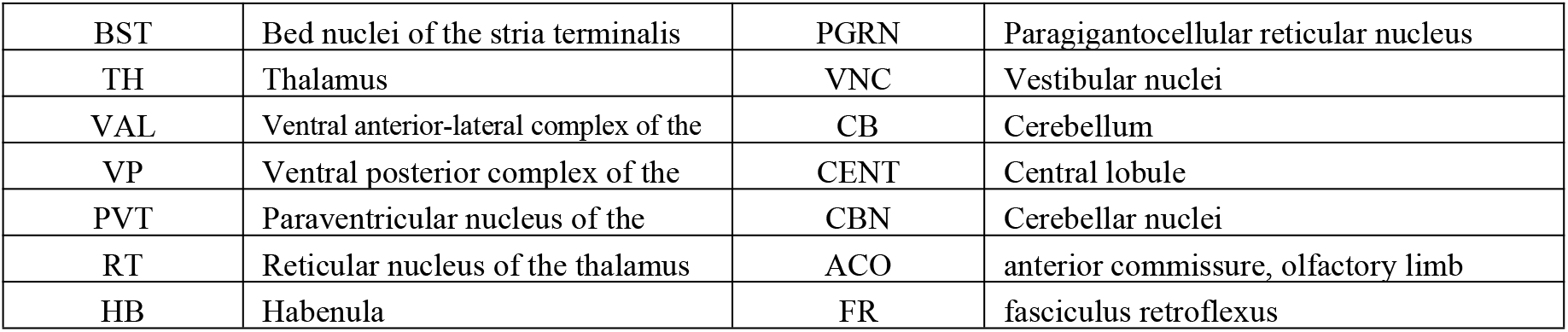
Abbreviations of anatomical structures of the mouse brain.

The distribution of RABV in brain after i.m., o.s. and i.n. infections is shown in Figures 2C, 2D and 2E, respectively. The infected regions regardless of inoculation route included the MO, BNST, SC, anteroventral periventricular nucleus (AVPV), accumbens nucleus (ACB), lateral septal nucleus (LS), paraventricular nucleus of hypothalamus (PVH), central amygdaloid nucleus (CEA), posterior hypothalamic area (PH), zona incerta (ZI), periaqueductal gray (PAG), and the hippocampal formation (HPF). In contrast, the infected regions associated with a particular inoculation route were the main olfactory bulb (MOB), dentate gyrus (DG), caudoputamen (CP), external globus pallidus (GPe), paraventricular nucleus of thalamus (PVT), lateral hypothalamic area (LH), vestibular nuclei (VNC), and the central lobule (CENT). I.m. inoculation resulted in that RABV primarily presented in the dorsolateral part of CP, but evenly distributed throughout the CP after o.s. inoculation. In contrast, i.n. inoculation resulted in almost no RABV infection in the CP (row 1, column 5 in Figure 2C-E, respectively). RABVs were distributed along the GPe and the gyrus in the cerebellum after i.m. inoculation, but not with o.s. or i.n. routes (row 2, column 1 and 8 in Figures 2C-E, respectively).

### 2.3. Quantification of RABV-infected cells in specific brain regions

To conduct a statistical analysis of the distribution of RABV in specific brain regions by different inoculation routes, we manually counted the number of EGFP-positive cells in the equidistant coronal sections of the cortex, striatum, pallidum, thalamus, hypothalamus, brain stem, and cerebellum, as well as EGFP-labeled cells per 100 μm^2^ to quantify the virus distribution in specific brain areas by using NIH ImageJ software (Figure 3). A majority of the cortical brain areas, except the CP and DG, showed a similar density of infection regardless of inoculation route. These areas included the paralemniscal nucleus (PL), somatosensory areas (SS), medial orbital (MO), and the infralimbic area (ILA). Similarly, the infection densities of the striatum (including accumbens nucleus (ACB), nucleus of the optic tract (OT), LS and CEA) and the hypothalamus (including PVH, AVPV, PH, lateral hypothalamic area (LHA) and ZI) were also similar among inoculation routes. In contrast, in the thalamus and brain stem, several differentially infected regions were observed, including the ventral anterior-lateral complex of the thalamus (VAL), interpeduncular nucleus (IPN), PVT, LH, and VNC.

**Figure.3.**
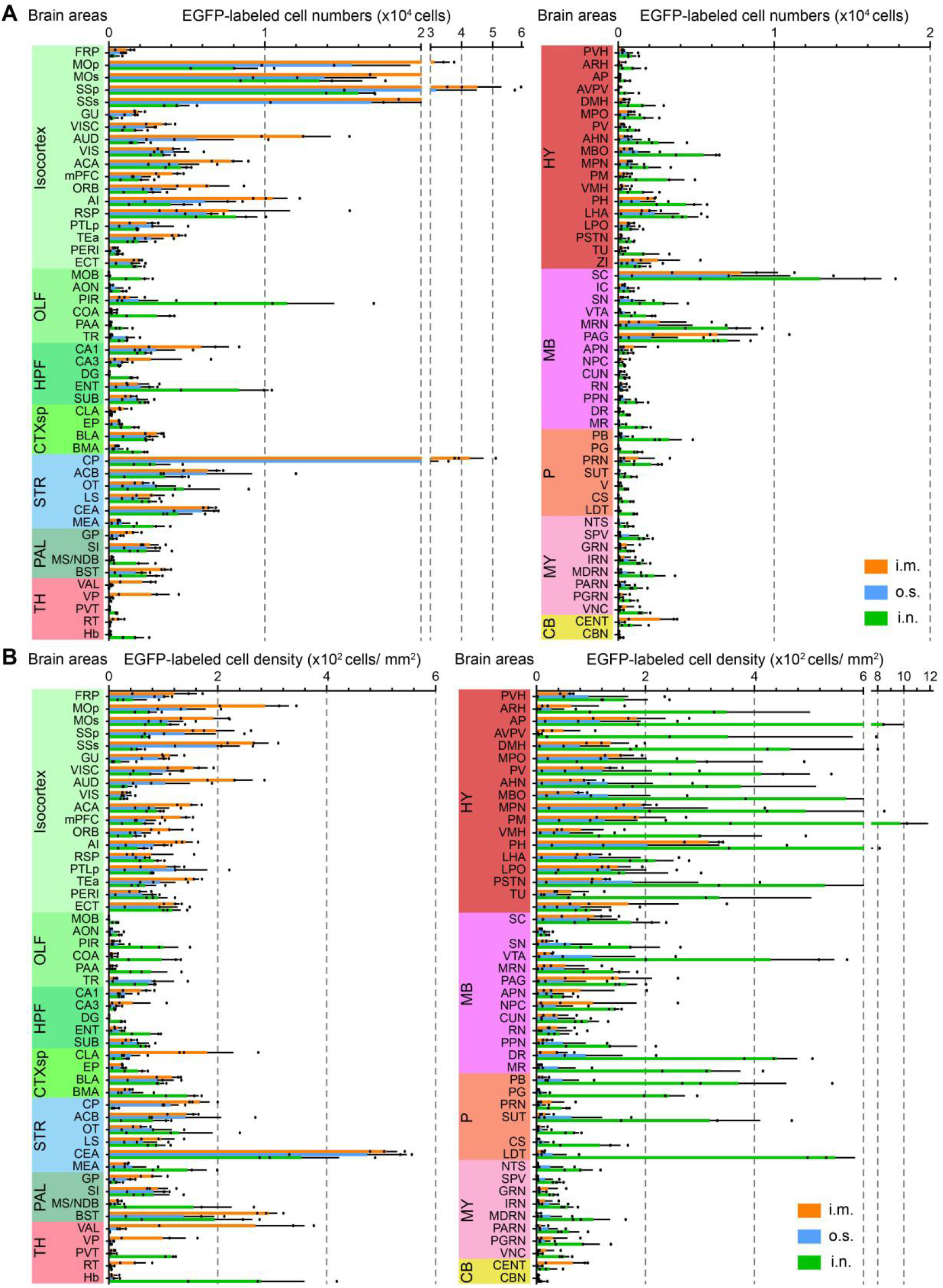
Comparison of RABV distribution in nuclei by different infection routes. (A-B) Total numbers and cell density of EGFP-labeled infected cells for each anatomical subregion, respectively. Data represent the mean ± SEM, and the abbreviations of anatomical subregions are listed in Supplementary Table 1.

The most remarkable characteristic of rabies patients is the arousal of fear. Several nuclei in the brain have been found to be related to fear, especially CEA, BST, HPF, PAG, PVH PL, ILA, ACB, LS and SC etc. ^14^. Interestingly, under all three infection routes, we identified the extensive RABV infection in these nuclei (Figure S5B) and the anatomical localization of these nuclei was shown in Figure S5A. Amygdala is known to play the central role in processing fear ^15, 16^, and is divided into cortical division and striatal division, the later one including medial amygdala (MEA) and central amygdala (CEA). Interestingly, we found that there is extensive distribution of RABV in CEA. Moreover, a newly found fear-related area, BST, as well as other fear-associated regions including HPF, PAG, PVH etc. were also extensively infected by RABV. Quantification of RABV-infected cells suggested that the infection routes had no significant impact on RABV distribution in the nuclei related to fear (Figure S5C), indicating that these nuclei are tightly associated with the pathogenesis of rabies.

### 2.4. Single-cell transcriptional profiling of RABV infection in the brain

Emerging evidences have shown that RABV is not a strict neurotropic virus and besides neurons it can infect other cell types ^17, 18^. To comprehensively investigate the infection susceptibility of diverse cell populations and their contributions to RABV pathogenesis, we performed droplet-based scRNA-seq (10×Genomics) on a total of six mouse brain samples including two from uninfected mice (healthy: n=2), two from mice with paralysis (paralyzed: n=2) and two from mice in the moribund stage post RABV infection (moribund: n=2) (Figure 4A). Following euthanasia, mouse brain tissue was obtained and rapidly digested to a single cell suspension. The single cell suspension from the uninfected brains was directly subjected to scRNA-seq. While the single cell suspension from RABV-infected brains was firstly enriched for cells containing EGFP signals (with active RABV infection) by FACS, and then analyzed by scRNA-seq (Figures 4A, S6A and S6B). With the unified single-cell analysis pipeline, ~0.77 billion unique transcripts were obtained from 54,452 cells. Among these cells, 16,199 cells (29.75%) were from the healthy mice, 13,417 cells (24.64%) were from the paralyzed mice and 24,836 cells (45.61%) were from the moribund mice (Figure S5C). All high-quality cells were integrated into an unbatched and comparable dataset and subjected to principal component analysis after correction for read depth and mitochondrial read counts (Figure S6D).

**Figure.4.**
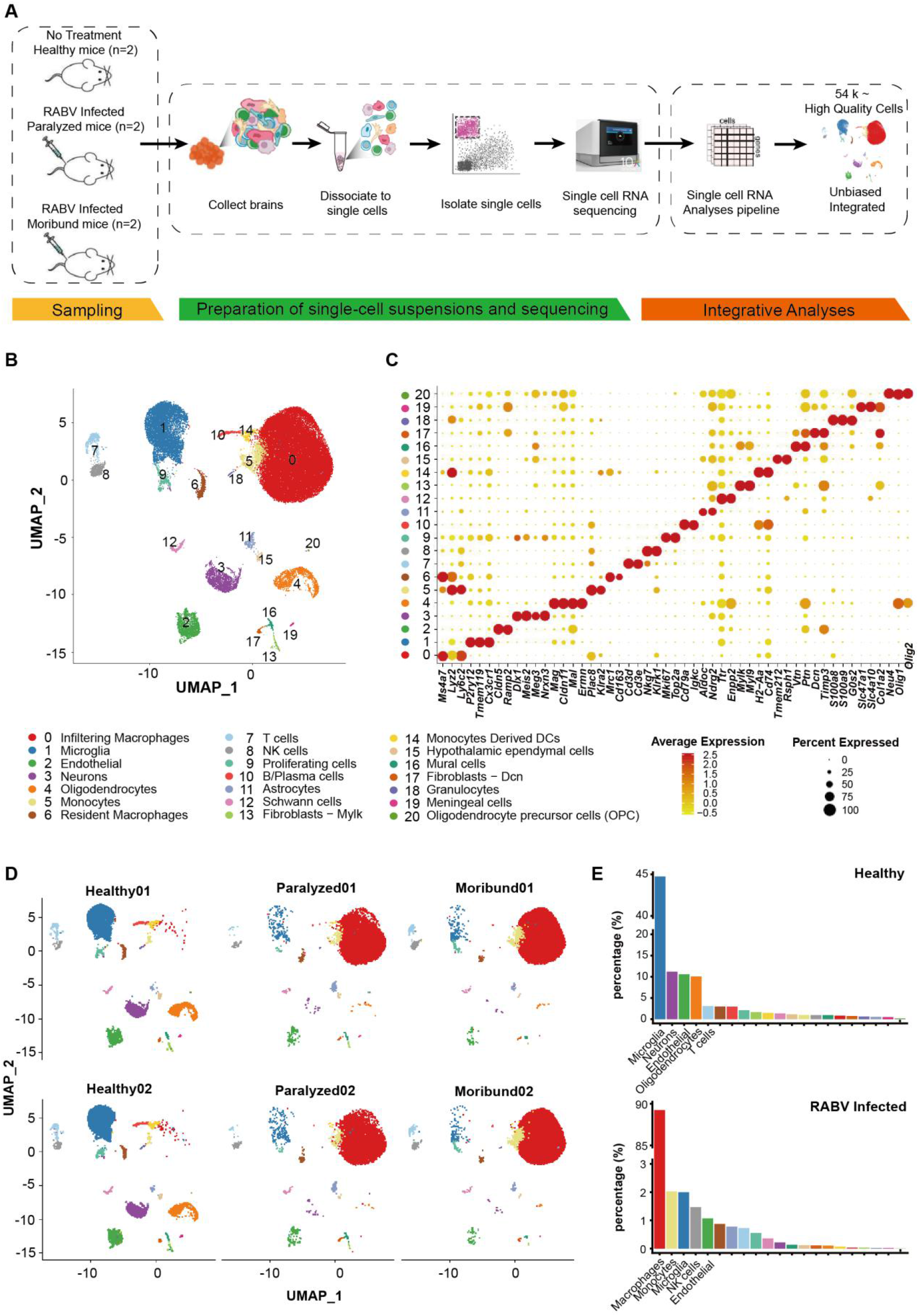
Single-cell transcriptional profiling of RABV infection in the brain. (A) A scheme showing the overall experimental design. scRNA-seq was applied to brain cells across the three conditions (healthy, paralyzed and moribund), and the output data were integrated and used for expression analyses. (B) Cellular populations identified. The UMAP projection of 54,452 single cells from healthy (n=2), paralyzed (n=2) and moribund (n=2) mouse brains, showing the formation of 21 clusters with the respective number and name labels. Each dot corresponds to a single cell, colored according to cell types. (C) Dotplot shows the expression level of canonical cell markers that used to identify clusters as represented in the UMAP plot. The color of dots represents expression levels and the size denotes expression percentage. (D) UMAP projection of each sample. Each dot corresponds to a single cell and is colored according to cell type. (E) Barplot shows the cell composition of healthy and the mean of the paralyzed and moribund conditions. Cellular populations were sorted in descending order and top five were labeled with cluster names.

Using graph-based clustering, uniform manifold approximation and projection (UMAP), we captured the transcriptomes of 21 high confidence cell clusters (Figure 4B) according to the expression of canonical markers (Figure 4C). Overall, the landscape contained the following cell lineages: T cells (*Cd3d*^+^), natural killing (NK) cells (*Klrk1*^+^), B/plasma cells (*Cd79a*^+^), monocytes (*Plac8*^+^), macrophages (*Ms4a7*^+^*Lyz2*^+^), granulocytes (*S100a8*^+^), monocytes derived dendritic cells (*H2-Aa*^+^), microglia (*P2ry12*^+^), neurons (*Meis2*^+^), oligodendrocytes (*Ermn*^+^) and its precursors (*Neu4*^+^), astrocytes (*Aldoc*^+^), ependymal cells (*Tmem212*^+^), schwann cells (*Ttr*^+^), endothelial cells (*Cldn5*^+^), fibroblasts (*Dcn*^+^), meningeal cells (*Slc4a10*^+^) and mural cells (*Vtn*^+^).

Consistent with previous studies ^19, 20^, the uninfected mouse brain consisted of microglia, neurons, endothelial, oligodendrocytes, and etc. (Figure 4D and Figure 4E above). We observed a great consistency in technical repeats within each sample condition, and the infected cells captured in the paralyzed and moribund condition have shown similar patterns (Figure 4D). After RABV infection, several cell types were significantly enriched, including macrophages, monocytes, microglia, and NK cells (Figure 4E below). To be noted, very few neurons were captured, possibly because neurons were very fragile post RABV infection and easy to be damaged during the processing of single-cell suspension and cell sorting by flow cytometry. To consolidate our scRNA-seq results, RABV infections in neurons, macrophages, microglia, and NK cells were confirmed by confocal microscopy (Figure S6E), virus titration (Figure S6F) and flow cytometry (Figures S6G and S6H) analysis.

### 2.5. Multiple roles of infiltrating macrophages during RABV infection in the brain

Macrophages are professional phagocytes that are integral to innate immune defense ^21^, and we observed a significant enrichment of macrophages (89.3%) after RABV infection (Figure 4E below). To gain an insight into their roles in neurotropic virus infection, 35,100 macrophages were obtained and re-clustered into four sub-clusters (Macro-C1 to C4) (Figures 5A, S7A and S7B). Macro-C1 and Macro-C2 were predominantly found in the paralyzed and moribund mouse brains, whereas Macro-C4 was predominantly observed in the healthy mouse brains. Macro-C3 could be found in both uninfected and infected brains, while its majority resided in the paralyzed and moribund mouse brains (Figures 5B and S7A).

**Figure.5.**
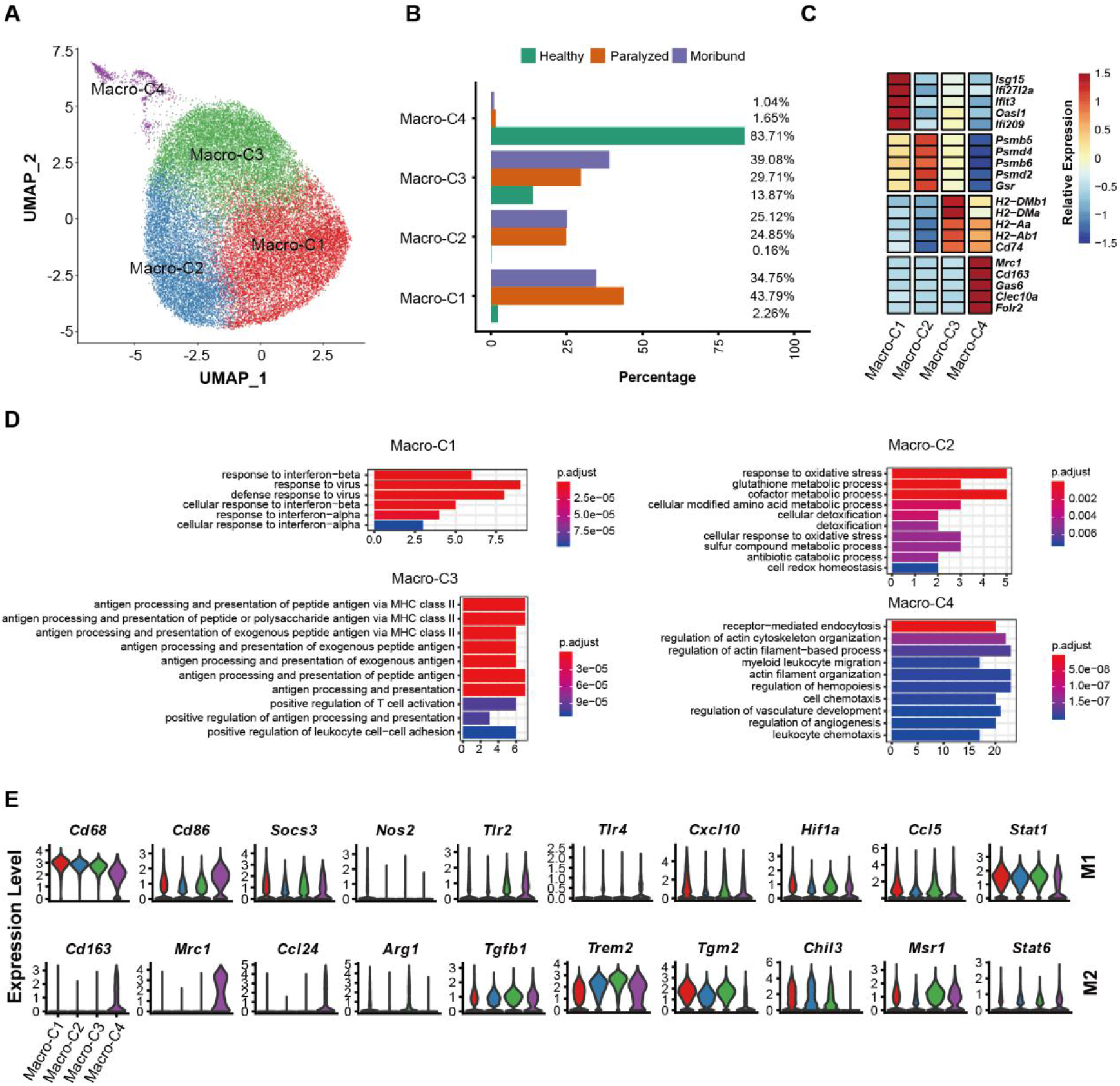
Transcriptomic features of macrophage subsets. (A) UMAP projection of 35,100 macrophages. Each dot corresponds to a single cell, colored according to subsets. (B) Barplot shows the condition preference of four macrophage clusters, colored according to three conditions. (C) Heatmap shows the relative expression levels of selected markers for four macrophage subsets. (D) Gene enrichment analyses of DEGs in four clusters. Gene Ontology (GO) terms are labeled with name and sorted by the adjusted p-value. Enriched gene counts were as the bar height and colored according to adjusted p-value. The top 10 enriched GO terms are shown. (E) Violin plots showing the expression distribution of M1 and M2 macrophage markers in four macrophage subsets.

To characterize each cluster, we first identified differentially expressed genes (DEGs) and then performed functional enrichment analyses (Figures 5C and 5D). Interferon response related DEGs, such as *Isg15, Ifi27l2a, Ifit3, Oasl1* and *Ifi209* were exclusively expressed in Macro-C1. In line with this, functional enrichment analyses revealed that Macro-C1 was an interferon response-related cluster. Macro-C2 was found to be related to cellular detoxification, metabolic and catabolic processes due to highly expressed proteases genes, such as *Psmb, Psmd* and *Gsr*. Macro-C3 highly expressed MHC-II genes, and functional enrichment analyses suggested that Macro-C3 might take part in antigen processing and presentation. Macro-C4 exclusively expressed *Mrc1, Cd163, Gas6, Clec10a, Folr2* and related to myeloid leukocyte migration, similar to the recently reported border-associated macrophages (BAMs), a brain resident macrophage residing in the dura mater, subdural meninges and choroid plexus 22.

Macrophages are conventionally classified into canonical M1 and M2 classes, the pro-inflammatory and anti-inflammatory macrophages, respectively ^23^. We found that no macrophage cluster exhibited only M1 or M2-like phenotype, whereas Macro-C4 exhibited a more M2-dominant gene signature, such as *Cd163, Mrc1* and *Ccl24* (Figure 5E). These data indicated that macrophage activation during RABV infection did not agree with the polarization model, either as discrete states or along a spectrum of alternative polarization trajectories.

Our data probably captured infiltrating macrophages asynchronously transitioning from one transcriptomic state to the next, we thus employed Monocle2 algorithm ^24^ to perform the pseudotime analysis. The inferred dynamic trajectory progressing exhibited a typical branched structure: with Macro-C1 as the root, Macro-C2 and Macro-C3 as the ending clusters (Figures 6A and S7C). To confirm that the ordering was correct, three marker genes were selected and plotted. As shown in Figures 6B and 6C, the expression level of *Isg15* decreased along the pseudotime; the expression level of *Psmb5* peaked in the middle of the pseudotime and the expression level of *H2-DMb1* was the highest at the end of the pseudotime, demonstrating a reasonable ordering.

**Figure.6.**
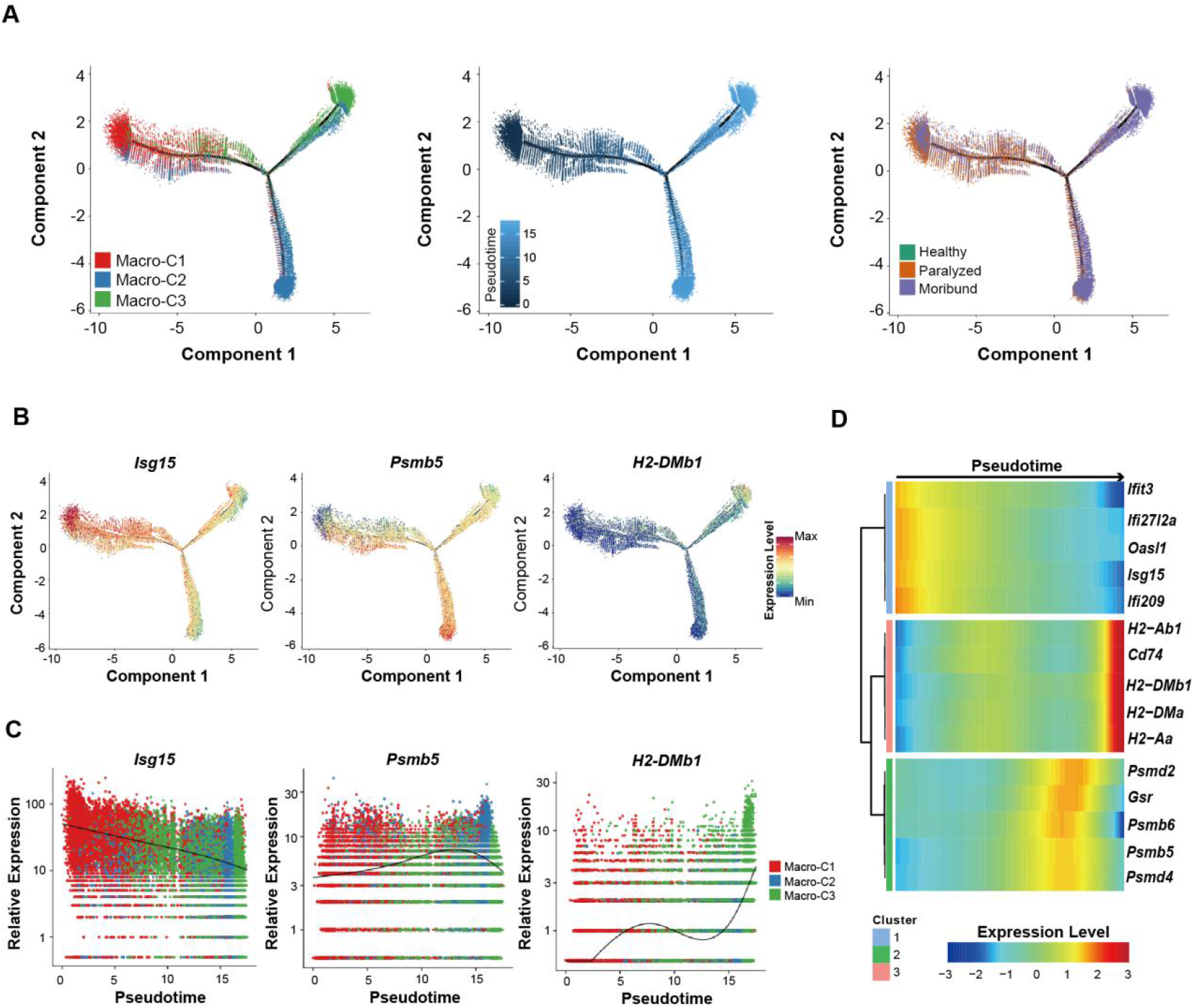
The potential developmental trajectory of three infiltrating macrophage subsets after RABV infection. (A) The potential developmental trajectory of three infiltrating macrophage subsets inferred by Monocle. Each dot corresponds to a single cell, colored according to subsets (left), pseudotime (middle) and conditions (right). (B) The expression levels of three exampled marker genes in the developmental trajectory. (C) Spline plots showing the expression kinetic trends of three exampled marker genes along the pseudotime. (D) Heatmap shows three modules of genes that co-vary across pseudotimes.

To understand the biological processes driving pseudotime components, we wondered which genes covary in expression with pseudotime. We clustered the representative genes identified as significantly covarying with pseudotime and identified three groups of genes expressed early, mid/mid–late and late (Figure 6D), consistent with the clusters identified above. Taken together, our results suggest that there are at least three different roles for the infiltrating macrophages, which we termed interferon-responsive macrophages, proteasome-active macrophages, and antigen processing and presentation macrophages during RABV infection in the brain.

### 2.6. RABV infection results in exhausted and apoptotic NK cells in the brain

NK cells are considered to be an important player of the innate immunity by controlling microbial infections ^25^. Intriguingly, we found that there was an obvious enrichment of NK cells in the mouse brain after RABV infection (Figure 4E). Re-clustering of the total of 719 NK cells revealed that there were at least three sub-clusters (NK-C1 to C3) (Figures 7A, S8A and S8B). The NK-C1 cluster was predominantly observed in the healthy brains, whereas sub-clusters NK-C2 and NK-C3 were enriched mostly in the brains from paralyzed and moribund mice (Figures 7B and S8A), respectively. To be noted, the NK-C1 cluster comprised highly expressed transcription factors and genes related to gene expression regulation and chromatin assembly. The NK-C2 cluster comprised highly expressed cytotoxic genes that take part in NK cell-mediated immunity, while the NK-C3 cluster was identified as an interferon response-related cluster (Figures 7C and 7D).

**Figure.7.**
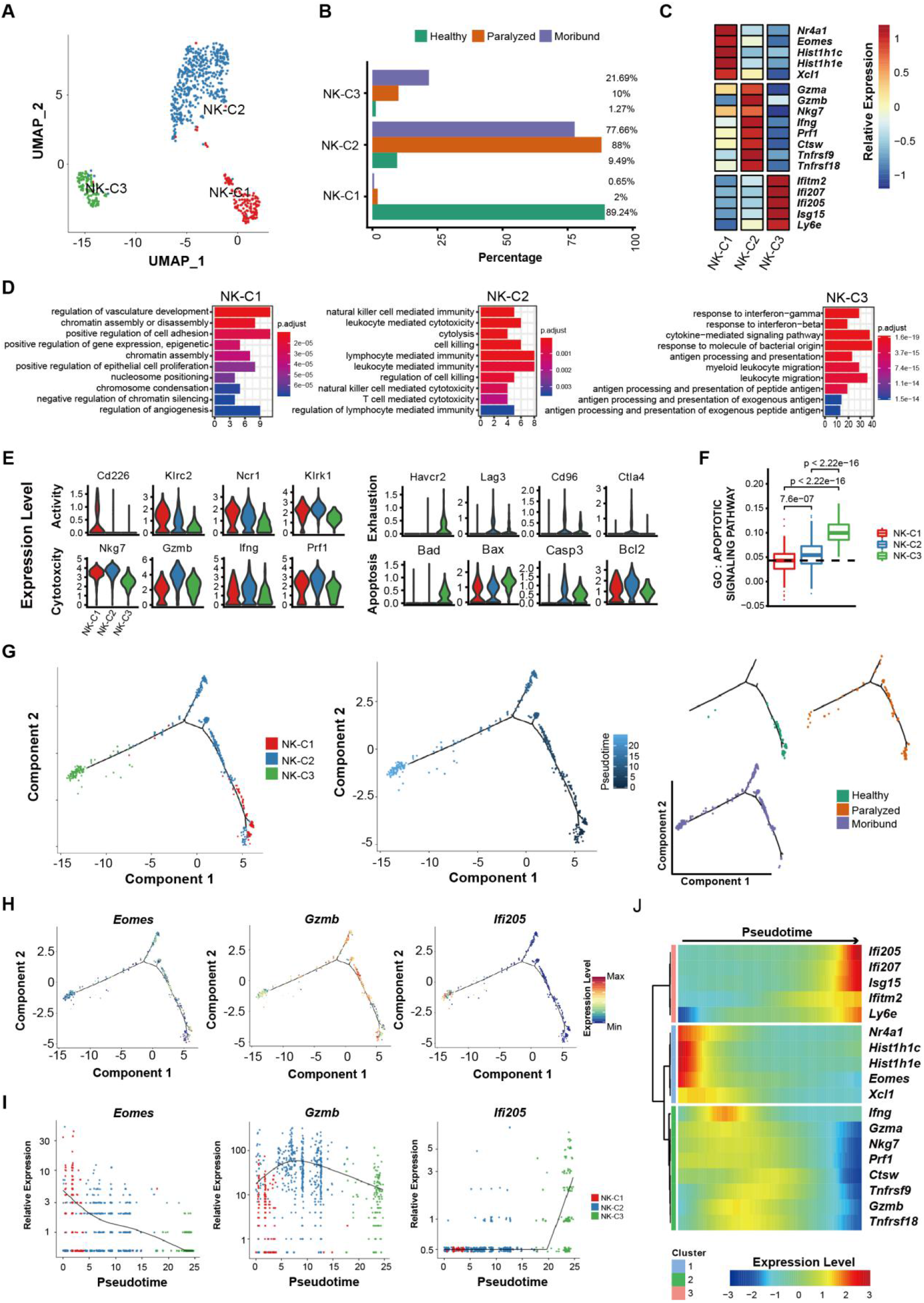
Transcriptomic features and potential developmental trajectory of NK cell subsets. (A) UMAP projection of 719 NK cells. Each dot corresponds to a single cell, colored according to subsets. (B) Barplot shows the condition preference of three NK subsets, colored according to three conditions. (C) Heatmap shows the relative expression levels of selected markers for three NK subsets. (D) Gene enrichment analyses of differentially expressed genes (DEGs) for three clusters. Gene Ontology (GO) terms are labeled with names and sorted by the adjusted p-value. Enriched gene count as bar height and is colored according to the adjusted p-value. The top 10 enriched GO terms are shown. (E) Violin plots showing the expression distribution of exampled markers in three NK subsets. (F) Box plots of the expression levels of GO apoptosis terms across different conditions. Conditions are shown in different colors. Horizontal lines represent median values, with whiskers extending to the farthest data point within a maximum of 1.5 × interquartile range. P-values were labeled and calculated by two-sided Wilcoxon rank sum test. (G) The potential developmental trajectory of three NK subsets. Each dot corresponds to a single cell, colored according to subsets (left), pseudotime (middle) and conditions (right). (H) The expression levels of three exampled marker genes in the developmental trajectory. (I) Spline plots showing the expression kinetic trends of three exampled marker genes along the pseudotime. (J) Heatmap shows three modules of genes that co-vary across pseudotimes.

NK cell functions are regulated by a balance between activating and inhibitory signals delivered by a multitude of receptors expressed on the cell surface ^25, 26^. The functional activated genes were down-regulated after RABV infection (e.g. *Cd226, Klrc2, Ncr1* and *Klrk1)*. In contrast, the exhausted genes were up-regulated after viral infection (e.g. *Havcr2, Lag3, Cd96* and *Ctla4*). The NK cell cytotoxic genes had the highest expression levels in the NK-C2 cluster, such as *Nkg7, Gzmb, Ifng* and *Prf1* (Figures 7E). The NK-C3 cluster showed the highest apoptotic degree across all three clusters (Figures 7E and 7F). These results indicate that NK cells were impaired by RABV infection with higher exhausted and apoptotic degree than that with healthy condition.

The inferred dynamic trajectory progressing of three NK cell types also exhibited a typical branched structure with NK-C1, NK-C2 and NK-C3 distributed as the root, middle and ending clusters, respectively. We found that the pseudotime was also consistent with developmental conditions where the start corresponded to the healthy condition and the end to the moribund condition (Figures 7G and S8C). The expression level of *Eomes* decreased along the pseudotime, the expression level of *Gzmb* peaked at the middle and of *Ifi205* was the highest at the end of the pseudotime (Figures 7H and 7I). To understand the biological processes driving pseudotime components, we further clustered the representative genes identified as significantly covarying with pseudotime and found the marker genes clustered by pseudotemporal expression pattern also revealed the same three clusters following similar kinetic trends (Figure 7J).

## 3. DISCUSSION

Due to the complexity of the brain, and lacking appropriate tools, researchers are facing great difficulty to investigate the invasion and distribution of neurotropic viruses in the brain, which impedes the invention of effective therapies for neurotropic viruses. The CLARITY, approach creates a tissue-hydrogel hybrid in which tissue components are replaced with exogenous elements. And then light-microscopy techniques can be used to access the entire mouse brain^27^. Recently, three-dimensional light sheet and confocal laser scanning microscopy was also applied to investigate the distribution of RABV in the brain and peripheral nerves^28^. fMOST technology makes it possible to map the spread of neurotropic viruses in the whole brain. It rapidly acquires a full-volume dataset in a whole mouse brain with resolution to a single neuron. The recently emerging scRNA-seq provides an efficient technique to investigate the transcriptomic characteristics during infection at single cell resolution. In this study, we investigated the spatial and cellular distribution of neurotropic virus infection in the mouse brain by jointly using fMOST and scRNA-seq.

We identified specific nuclei in which viral spread or the viral load depended on the route of infection. Considering the role infection route has on the distribution of RABV in the brain, those regions commonly infected are likely critical for deciphering RABV pathogenesis and the potential targets for future therapeutics. Considering the influence of infection route on viral distribution, knowledge of those regions that are commonly infected is especially important for understanding RABV pathogenesis. Previous studies have suggested that aggressive behavior is critical for RABV pathogenesis, because it leads to the efficient transmission of RABV to other hosts by the rabid animals. In RABV-infected skunks, the red nucleus (RN) and midbrain raphe nuclei were found to be associated with aggressive behavior ^29^. Notably, our data indicate that RABV is distributed to the RN only post i.m. inoculation (Figure 3).

Fear is another cardinal characteristic of rabies, which is an emotion that has powerful influence on behavior and physiology. Fears, depending on the type of threat stimuli, are processed in independent neural circuits that involve the amygdala and downstream hypothalamic and brainstem circuits ^16^. Rabies patients often have abnormal symptoms like hydrophobia, aerophobia and phonophobia indicating that there is a disorder in the processing of fear. In this study, we found that several nuclei related to fear, such as CEA, BNST, PAG, and BLA, were extensively infected by RABV (Figure S4). Notably, the route of infection made no difference in the viral load in these areas, indicating that RABV infection in fear-related areas is independent of the infection route.

Besides fear, inspiratory muscle spasms are another cardinal feature of rabies infection. In the late stages, periodic and ataxic breathing is often observed ^30^. Previous studies have shown that the midbrain and medulla, especially the pontine tegmentum, are the regions most affected by RABV-induced inflammation ^30^. Accumulating evidence suggest that pontine nuclei, including Kölliker–Fuse (K-F) and parabrachial (PB) nuclei comprise the pontine respiratory group (PRG), which regulates the inspiratory-expiratory phase transition ^31, 32^. Other studies report that neurons in the superior colliculus (SC) ^33^, RN ^33^, and intermediate reticular nucleus (IRN) ^34^ also influence the respiratory cycle. In our study we showed that the PB, SC, RN and IRN were extensively infected by RABV regardless of the infection route (Figure 3), suggesting that the abnormal breathing may be related to the neuronal damage caused by RABV infection in these nuclei.

The fMOST results provide the spacious distribution of RABV in the brain, but could not determine the exact infected cell types. To further investigate the cellular distribution of RABV, scRNA-seq analysis was implemented and the results revealed that cell types in the brain from healthy mice consisted of microglia, neurons, endothelial, oligodendrocytes, and etc. which was consistent with the previous reports ^19, 20^. Different from fMOST data which showed that most of the RABV-infected cells were neurons, the vast majority of RABV-positive cells isolated from the brains of paralyzed and moribund mice were immune cells, such as macrophages and NK cells. However, the proportion of RABV-positive neurons among single cell suspension was relatively low. A possible reason for this paradox is that neurons are hard to pass through the cell sorting by flow cytometry due to its morphology and size according to the previous studies ^35, 36^. Another reason is because neurons are quite fragile, especially post RABV infection, and most neurons are damaged or even died during the processing of preparing single cell suspension, which involves repeated enzyme digestion and rotation. Before scRNA-seq analysis, we enriched EGFP-positive cells by flow cytometry and for the first time we identified that RABV could cause infection in microglia /macrophages and NK cells in the mouse brain.

Monocytes are the first line in defending against viral infection. Due to its short half-life, it is difficult for virus to replicate in monocytes. Upon infection, monocytes change their cytokine/chemokine pattern and thus differentiate into long-lived macrophages ^37^. Macrophages play a critical role in viral infection: they can restrict viral infection, while viruses can utilize macrophages as vessels for persistence or dissemination in tissues ^37^. Macrophages in the brain include resident macrophages (microglia) and monocytes derived macrophages that infiltrate from the periphery post pathogen infection. CNS resident macrophages comprise microglia and border-associated macrophages (BAMs) ^38^. We mainly observed the brain resident macrophage (Macro-C4) in healthy mice, while the macrophages in RABV-infected groups were mainly belonging to the functional macrophage types Macro-C1, Macro-C2 and Macro-C3. This is consistent with a previous study which found that peripheral monocytes quickly infiltrated into the CNS after viral infection, while resident macrophages were redundant for antigen presentation and underwent apoptosis during the chronic phase ^39^.

In our inferred developmental trajectory we showed that interferon responses cluster (Macro-C1) appeared earlier than cellular detoxification cluster (Macro-C2) and antigen processing and presentation cluster (Macro-C3), which is consistent with the fact that interferon induces the polarization of macrophages following virus infection, and then macrophages participate in early immune response through phagocytosis and antigen presentation ^40^. It is also noteworthy that understanding of macrophages’ role in neurotropic virus infection focused on their phagocytosis and antigen presentation. However, recent studies have found that macrophages might act as a virus reservoir during infection with various viruses, assist in virus replication or long-term existence in the body, and bring viruses into other cells, even the CNS ^37^. Virus reservoir macrophages are neither M1 nor M2 type, but exist in the form of M1 / M2 continuum. They usually show abolished apoptosis and restricted cytopathic effects, which facilitates the virus production ^37^. In our study, we also found a group of such kind cells, suggesting that macrophages not only perform phagocytosis and antigen presentation during RABV infection, but also may play a role in assisting the virus to escape the immune response.

NK cells can eliminate infected cells by recognizing cell surface receptors or secreting killing media like perforin or TNF_α_. However, growing evidences showed that various viruses can induce apoptosis and lead to the exhaustion of NK cells ^41, 42^. Interestingly, our inferred developmental trajectory of three NK subsets exhibited a branched structure with NK-C1, NK-C2 and NK-C3 distributed as the root, middle and ending clusters, respectively, reflecting the transition process of NK cells following virus infection from the normal state to the killing state and then to the apoptotic state. The higher apoptotic status in NK-C2 and NK-C3 than NK-C1 may indicate that RABV may escape the innate immune response by inducing the apoptosis of NK cells. In a previous study, it was found that RABV could induce the apoptosis of infiltrating T cells by upregulating of FasL and B7-H1 in the surface of infected neurons, and then evaded host T cell defenses ^43^. Thus we speculate that RABV can employ the same strategy to evade the killing by NK cells.

In summary, we utilized fMOST technology to reveal the susceptible nuclei infected by RABV in the mouse brain, and take advantage of scRNA-seq technology to analyze the RABV-infected cells and illustrate the roles of some immune cells. The joint use of these two technologies allowed us to portray an integrated map of RABV infection in the mouse brain. Our results shed a light for future investigation of the pathogenesis and clinical therapy of rabies and other neurotropic viruses such as ZIKA and dengue viruses.

## 4. Experimental Section

### Ethics statement

The experiments involving mice in this study were performed in accordance with the recommendations in the Guide for the Care and Use of Laboratory Animals of the Ministry of Science and Technology of China and were approved by the Scientific Ethics Committee of Huazhong Agricultural University (permit number HZAUMO-2016-052).

### Brain sample preparation for imaging

Groups of six-week-old female C57BL/6 mice were inoculated with 10×LD_50_ RABV-EGFP by intramuscular (i.m.), otic subcutaneous (o.s.), or intranasal (i.n.) route (n=3). In severe paralysis, mice were anesthetized with ketamine/xylazine and then perfused by intra-cardiac injection of PBS followed by 10% neutral-buffered formalin. Brains were removed and fixed with 10% neutral-buffered formalin for 24 hours at 4 °C. After fixation, each brain was rinsed overnight at 4 °C with PBS and embedded with oxidized agarose before imaging.

### Imaging by fMOST

The agarose-embedded brains were imaged by fMOST system^8^. During the process of imaging, the brain was immersed in water and a water immersion objective (1.0 NA, 20×) was used. The fluorescent signals were detected via a scientific complementary metaloxide semiconductor (sCMOS) camera, a modern scientific camera with highly sensitive and high-speed. A 3D stage accurately moved the brain mosaic-by-mosaic to extend the field of view in the imaging part, and then the stage moved the brain towards the oscillatory blade to remove the imaged tissue in the vibrating sectioning part. The full-volumetric imaging was performed with the cycle of imaging and sectioning until the whole-brain dataset was finished. For a single mouse brain, the dataset included approximately 3000 coronal sections was collected in three days at a voxel resolution of 0.32 μm × 0.32 μm ×5 μm.

### Image reconstruction and analysis

Image preprocessing for mosaic stitching and uneven lateral illumination correction was performed as reported previously^7^. We visualized the datasets using Amira software (v 5.2.2, FEI, France) to generate the figures of maximum intensity projection, volume, surface rendering, and for the movies. The preprocessed dataset was imported into Amira software using a desktop graphical workstation (T7600, Dell Inc., USA).

### Quantification of RABV-infected cells

To quantify the infected cells, the EGFP-labeled cells were automatically segmented and coregistered to Mouse Reference Atlas as previously described^44, 45^. Briefly, each coronal section was background subtracted, Gaussian filtered, and threshold segmented to binary image, and the infected cells were segmented with individually adjusted binary thresholds according to varying fluorescent intensities. And then the soma coordinates were warped and coregistered to the corresponding Allen Mouse Reference Atlas (Allen Institute for Brain Science) coordinate using non-rigid registration with free-form deformation. The anatomical sub-regions of which infected cells belonged to were then mapped to the Allen Mouse Reference Atlas. We calculated the total number and the density of EGFP-labeled infected cells for each anatomical sub-region with different infection routes (n = 3 per infection route).

### Cell sorting and scRNA-seq

Groups of six-week-old female C57BL/6 mice were i.m. inoculated with 10×LD_50_ RABV-EGFP. At the stage of paralysis and moribund, mice were anesthetized with ketamine/xylazine and then brains were collected. Single-cell suspensions were obtained with adult brain dissociation kit (Miltenyi Biotec, 130-107-677). EGFP positive cells were sorted using the GFP channels of the Bio-Rad S3e instrument. cDNA libraries were prepared from single-cell suspensions following the instruction of 10×Genomics 3’ V3. Cells obtained from the whole brain of the healthy mice with the same method and enriched by flow cytometry were used as control. RNA-sequencing was performed by Novogene (Nanjing, China).

### scRNA-seq data processing

Raw gene expression matrices were generated for each sample by the Cell Ranger (Version 3.0.2) Pipeline coupled with mouse reference version mm10. The output filtered gene expression matrices were analyzed by R software (Version 3.5.3) with the Seurat 46 package (Version 3.0.0). In brief, genes expressed at a proportion > 0.1% of the data and cells with > 200 genes detected were selected for further analyses. Low-quality cells were removed if they met the following criteria: 1) < 500 or > 70,000 UMIs; 2) < 200 or > 7,500 genes; or 3) > 10% UMIs derived from the mitochondrial genome. After removal of low-quality cells, the gene expression matrices were normalized by the *NormalizeData* function, and 2000 features with high cell-to-cell variation were calculated using the *FindVariableFeatures* function. To reduce dimensionality of the datasets, the *RunPCA* function was conducted with default parameters on linear-transformation scaled data generated by the *ScaleData* function. Next, the *ElbowPlot, DimHeatmap* and *JackStrawPlot* functions were used to identify the true dimensionality of each dataset, as recommended by the Seurat developers. Finally, we clustered cells using the *FindNeighbors* and *FindClusters* functions, and performed non-linear dimensional reduction with the *RunUMAP* function with default settings. All details regarding the Seurat analyses performed in this work can be found in the website tutorial (https://satijalab.org/seurat/v3.0/pbmc3k_tutorial.html).

### Multiple dataset integration

To compare cell types and proportions across three conditions, we employed the integration methods described at https://satijalab.org/seurat/v3.0/integration.html47. The Seurat package (Version 3.0.0) was used to assemble multiple distinct scRNA-seq datasets into an integrated and unbatched dataset. In brief, we identified 2000 features with high cell-to-cell variation as described above. Secondly, we identified “anchors” between individual datasets with the *FindIntegrationAnchors* function and inputted these “anchors” into the *IntegrateData* function to create a “batch-corrected” expression matrix of all cells, which allowed cells from different datasets to be integrated and analyzed together.

### Sub-clustering of macrophages and NK cells

First, macrophages were extracted from the overview integrated dataset. Next, the major cell types were integrated for further sub-clustering. After integration, genes were scaled to unit variance. Scaling, PCA and clustering were performed as described above. NK cells were also extracted and sub-clustered using the procedure used for macrophages.

### Cell type annotation and cluster markers identification

After non-linear dimensional reduction and projection of all cells into two-dimensional space by UMAP, cells clustered together according to common features. The *FindAllMarkers* function in Seurat was used to find markers for each of the identified clusters with “wilcox” method and obey “min.pct = 0.2” and “logfc.threshold = 0.25”. Clusters were then classified and annotated based on expressions of canonical markers of particular cell types. Clusters expressing two or more canonical cell-type markers were classified as doublet cells and excluded from further analysis.

### Differentially expressed genes (DEGs) identification and functional enrichment

Differential gene expression testing was performed using the *FindMarkers* function in Seurat, and the Benjamini-Hochberg method was used to estimate the false discovery rate (FDR). DEGs were filtered using a minimum fold change of 1.5 and a maximum FDR value of 0.01. Enrichment analysis for the functions of the DEGs was conducted using clusterProfiler 48 in default parameters and the Benjamini-Hochberg method was used to estimate FDR. Gene sets were derived from the Biological Process of GO Ontology.

### Defining cell state scores

We used cell scores in order to evaluate the degree to which individual cells expressed a certain pre-defined expressed gene set. The cell scores were initially based on the average expression of the genes from the pre-defined gene set in the respective cell ^49^. The *AddModuleScore* function in Seurat was used to implement the method with default settings. We used APOPTOTIC SIGNALING PATHWAY (GO: 0097190) to define the apoptosis score.

### Pseudotime trajectory inference

We applied the Monocle^24^(version 2) algorithm to determine the potential lineage differentiation between diverse cell populations refer to the tutorial here (http://cole-trapnell-lab.github.io/monocle-release/docs/). First, store data in newCellDataSet object with the parameter “expressionFamily= negbinomial.size()” and “lowerDetectionLimit = 0.5”. Before construct single cell trajectories, size factors and dispersions were estimated and filtered low-quality cells with default settings. Then the trajectory was inferred with the default parameters of Monocle after dimension reduction and cell ordering based on top 500 genes differing between clusters. Finally, the results of inferred pseudotime trajectory were presented and shown with the first two components.

### Statistical analyses

The statistical tools, methods and thresholds for each analysis are explicitly described with the results or detailed in the Figure Legends or Methods sections.

## Supporting Information

Document S1. Supplemental data

## Data and Code Availability

The raw sequence data reported in this paper have been deposited in the Genome Sequence Archive at the BIG Data Center, Beijing Institute of Genomics (BIG), Chinese Academy of Sciences, under the accession number CRA002563 and are publicly accessible at http://bigd.big.ac.cn/gsa. Custom scripts for analyzing data are available upon reasonable request.

## Acknowledgements

We thank the Optical Bioimaging Core Facility of HUST for support with data acquisition. This study was partially supported by Guangdong Major Project of Basic and Applied Basic Research (2020B0301030007) and the National Natural Science Foundation of China (No. 31872451 to L.Z.; No. 31722003 and No.31770925 to F.B.; No. 21778020 to H.Y.H.). This work was also supported partially by the Science Fund for Creative Research Group of China (61721092), NSFC Grants (81827901, 91749209), and the Director Fund of WNLO. to H.G..

## Conflict of Interest

The authors declare no competing interests.

## Author Contributions

Conceptualization, L.Z., F.B., H.G.; Investigation, Y.C.Z., X.D.X., B.L., Y.D.C., S.M.H., X.N.L., Y. L. Y., D. Y. T., Z. C. L., B. K. S., W. L., L. L, Q. W, J. X. D., M. Z., Z. F. F.; Visualization, B. L., Y. L. Y.; Data analyses, X.D.X., B.L., Y.C.Z.; Funding Acquisition, L.Z., F.B., H.G., H.Y.H.; All authors read and approved the final manuscript.

## Notes

### Competing Interest Statement

The authors have declared no competing interest.

