## Supplementary data for "A spatial and cellular distribution of neurotropic virus infection in the mouse brain revealed by fMOST and single cell RNA-seq"

### Supplementary Experimental Procedures

#### Cells, Antibodies, and Mice

BSR cells, a cloned cell line derived from BHK-21 cells, and neuroblastoma (NA) cells were cultured in Dulbecco's modified Eagle's medium (DMEM) (Mediatech, Herndon, VA) supplemented with 10% fetal bovine serum (FBS) (Gibco, Grand Island, NY). Fluorescein isothiocyanate (FITC)-conjugated antibodies against RABV-N protein were purchased from Fujirebio Diagnostics, Inc. (Malvern, PA). NKp30 mouse monoclonal antibody (Santa Cruz, sc-33647); RABV-P rabbit polyclonal antibody (prepared in our own lab); Alexa Fluor 488-conjugated goat anti-mouse antibody (Invitrogen, R37120); Alexa Fluor 594-conjugated anti-rabbit antibody (Invitrogen, A11012); DAPI (Invitrogen, D1306) were used in confocal microscopy. Anti-IBAI antibody (ab5076) was used to identify microglia by immunofluorescence. Three-day-old and six-week-old female C57BL/6 mice were purchased from the Hubei Center for Disease Control, Wuhan, China and handled according to protocols approved by the Scientific Ethics Committee of Huazhong Agricultural University (permit number: HZAUMO-2016-052). The mouse experiments associated with RABV infection were all operated in the BSL-2 laboratory in Huazhong Agricultural University.

#### Construction of RABV Expressing EGFP

The recombinant RABV CVS-B2c was constructed as previously described [1]. A transcription unit containing the BsiWI and NheI restriction sites was introduced between the G- and L-coding sequences by deleting the pseudogene. EGFP was cloned and inserted between the BsiWI and NheI restriction sites, resulting in RABV-EGFP. The full length clone of RABV-EGFP and four helper plasmids that expressed the N, P, G and L genes were separately transfected into BSR cells using the SuperFect transfection reagent (Qiagen, Valencia, CA) according to procedures previously described [1]. Rescued RABV-EGFP was detected with FITC-conjugated antibodies against RABV-N under an Olympus IX51 fluorescence microscope.

#### Virus Titration

Virus titration was determined as described previously[2]. Briefly, 10-fold serial dilutions of virus were incubated with BSR cells in 96-well plates at 34 °C for 48 h. Cells were then fixed with 80%

ice-cold acetone for 15 min and stained with FITC-conjugated anti-RABV-N antibody at 37 °C for 45 min. Antigen-positive foci were counted under an Olympus IX51 fluorescence microscope, and virus titers were calculated using the Reed-Muench formula and presented as fluorescent focus units per milliliter (FFU/mL) [3]. All titrations were carried out in quadruplicate.

**Pathogenicity of RABV in Mice**

Two groups of 6-week old female C57BL/6 mice (n=10) were inoculated with $5\times10^5$ FFU RABV or RABV-EGFP by i.m. injection. Mice were monitored daily for clinical signs of disease for 3 weeks. The clinical signs were scored using a scale of 0 to 5 as described previously [4]: 0, no clinical signs; 1, disordered movement; 2, ruffled fur, hunched back; 3, trembling and shaking; 4, complete paralysis; 5, death. Mice that lost more than 25% body weight were euthanized with $CO_2$ and the survivor ratio was calculated.

**Quantitative Reverse Transcription-PCR (qPCR)**

Viral load in the mouse brain was quantitated by qPCR as described previously[5]. Total RNA was isolated from different sections of the brain using TRIzol reagent (Invitrogen), and then reverse transcribed using ReverTra Ace qPCR RT Master Mix (Toyobo, FSQ-201). qPCR was performed using SYBR Green Supermix (Bio-Rad) according to manufacturer's protocol. A standard curve was generated from serially diluted pcDNA3.1-N. RABV N mRNA copy numbers were normalized to 1 g of total RNA. Primer pairs used for amplification of RABV N mRNA: For, 5'-GATCGTGGAACACCATACCC-3'; Rev, 5'-TTCATAAGCGGTGACGACTG -3'.

**Frozen Sections**

RABV-infected mice were euthanized when they lost around 25% of their body weight. Their brains were harvested and fixed in 4% neutral buffered paraformaldehyde (PFA). The brain was immersed in 20 ml 30% sucrose for 48-72 h and then coated with frozen-section embedding compound (OCT, Fisher Scientific, PA). After briefly drying, the brains were sectioned using a Leica frozen slicer. IFA staining in the samples was observed under an Olympus IX51 microscope.

**Determination of $LD_{50}$ of RABV-EGFP**

Three groups of six-week-old female C57BL/6 mice (n=10) were inoculated with RABV-EGFP by the i.m. (100 μl /mice), the o.s. (30 μl /mice), or the i.n. route (10 μl /mice). Virus was diluted to the appropriate concentration with DMEM. The clinical signs and survivor numbers were recorded daily for 3 weeks as described previously [2]. When mice became moribund they were euthanized by $CO_2$. The fifty percent lethal dose ($LD_{50}$) for each route of infection was calculated as described by Reed and Muench [3].

**Flow cytometry**

Groups of C57BL/6 mice inoculated with RABV were euthanized at the stage of moribund, and brains were collected and dissociated into single cells. To select RABV positive macrophages, cells were stained with FITC anti-RABV-N, PE anti-mouse CD11b (clone M1/70) and PE-Cy5.5 anti-mouse CD45 (clone 30-F11). RABV-$P^+CD11b^+CD45^+$ cells were defined as RABV positive macrophage. To select RABV positive NK cells, cells were stained with FITC anti-RABV-P, PE-Cy7 anti-mouse CD3 (clone 17A2) and APC anti-mouse NK1.1 (clone PK136). RABV-$P^+CD3^-NK1.1^+$ cells were then defined as RABV positive NK cells.

**Isolation and culture of primary cells**

(1) Microglia: three-day-old C57BL/6 mice were euthanized by $CO_2$, and brains was removed, detached from the meninges, and shredded into small pieces around 1-2 $mm^3$. Then the tissues were digested with trypsin and DNase and centrifuge. Cell pellet was resuspended after centrifugation with 10% serum medium and cells were plated for cultivation. After 8 days, generation of obvious astrocytes and microglia can be observed under a microscope. Microglial cells are easy to shake off, while primary astrocytes are closely attached to plates.

(2) Bone marrow-derived macrophages: six-week-old C57BL/6 mice were euthanized by $CO_2$. Hind leg muscles were removed to obtain thighs. Then both ends of the thighs were cut and bone marrows were blown into serum-free medium. Cells were cultured with a medium containing 10% FBS and 30% L929 cell supernatant for 7 days, and then macrophages can be harvested for further studies.

**Supplementary Figures and Figure Legends**

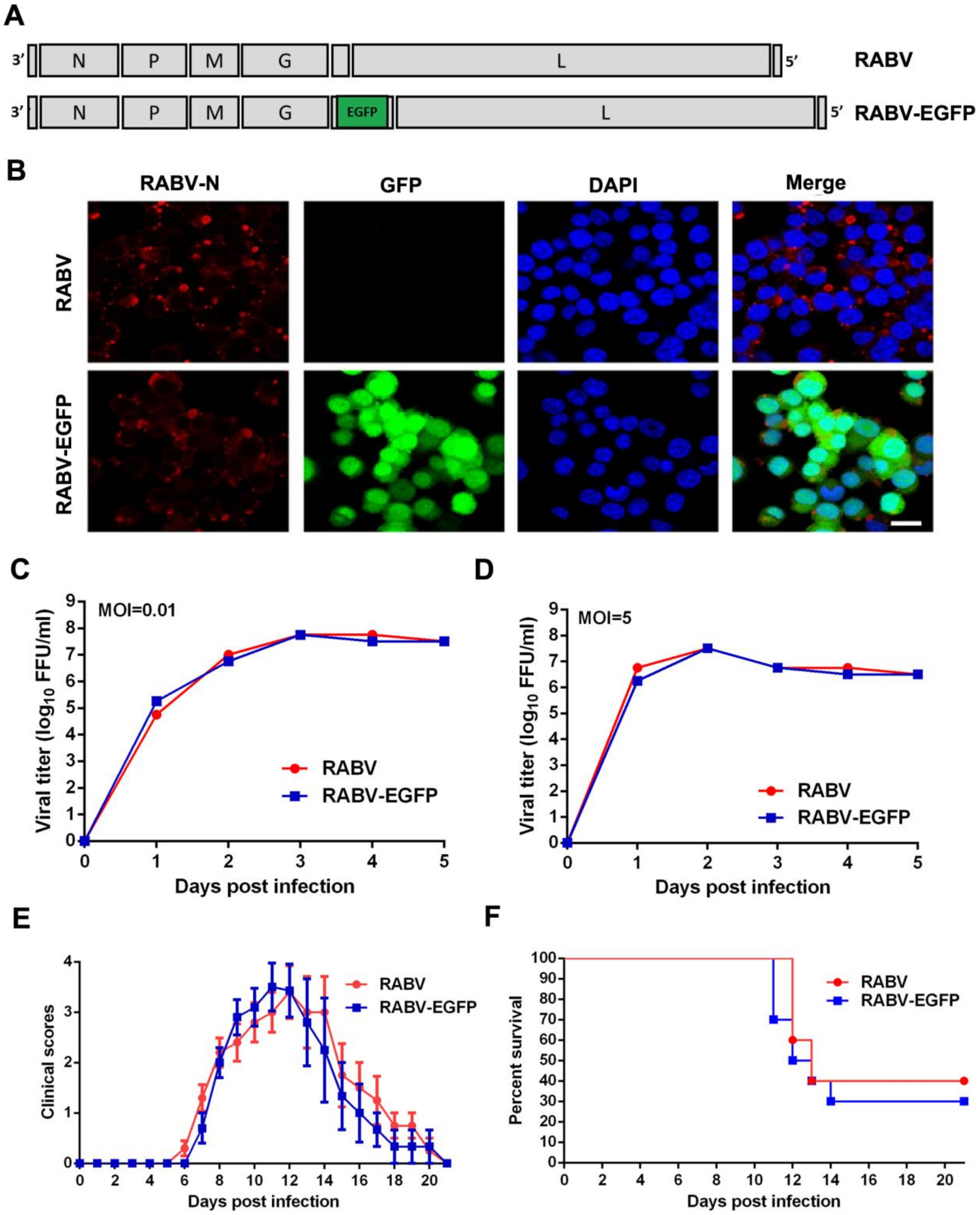


**Additional file 1: Figure S1 Construction and characterization of the recombinant RABV expressing EGFP (RABV-EGFP), Related to Figure 1**. (A-B) Strategy for the construction of RABV-EGFP is shown in panel A. EGFP was inserted into the genome of the recombinant RABV strain CVS-B2c between the G and L genes, and RABV-EGFP was rescued and confirmed in NA cells (B). (C-D) The growth kinetics of RABV and RABV-EGFP in BSR cells. BSR cells were infected with RABV or RABV-EGFP at MOI=0.01 and 5. Virus titers in the cell supernatants were determined by direct immunofluorescence (IFA). Three technical replicates were completed for each sample. (E-F) Two groups of female C57BL/6 mice (6-8-week-old, n=10) were inoculated i.m. with $6\times10^4$ FFU RABV or RABV-EGFP. The mice were monitored daily for three weeks and clinical scores (E) and the survivor ratios (F) were recorded.

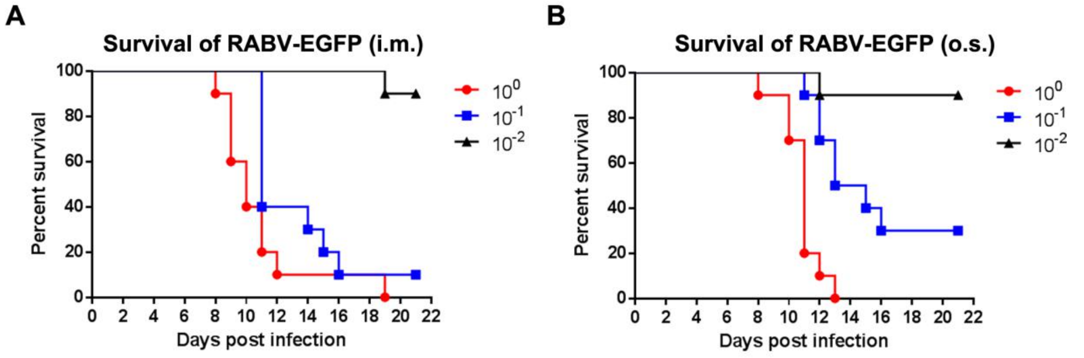


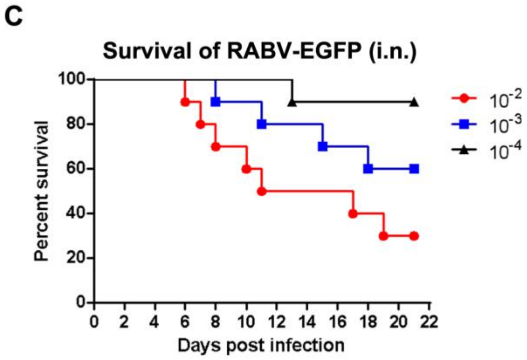


D

**$LD_{50}$ of RABV-EGFP**

| Route of Infection | $LD_{50}$ |
|---|---|
| i.m. | **$10^{-1.50}$/0.1 ml** |
| o.s. | **$10^{-1.35}$/0.03 ml** |
| i.n. | **$10^{-2.68}$/0.02 ml** |

**Additional file 2: Figure S2 Determination of RABV-EGFP $LD_{50}$ by different infection routes, Related to Figure 1**. Groups of female C57BL /6 mice (6-8-week-old, n=10) were inoculated with different dilutions of RABV-EGFP by i.m. (100 µl /mouse), o.s. (30 µl /mouse), or i.n. (20 µl /mouse). (A-C) Mice were monitored daily for three weeks and the survivor ratios were recorded. (D) $LD_{50}$ was calculated according to Reed and Muench formula.

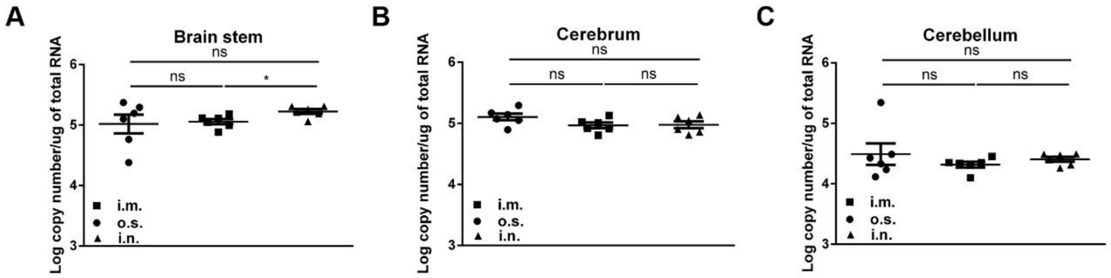


**Additional file 3: Figure S3 Viral load of RABV-EGFP in the mouse brain, Related to Figure 1.** Three groups of female C57BL /6 mice (6-8-week-old, n=6) were inoculated with 10×$LD_{50}$ of RABV-EGFP by i.m., o.s., or i.n. When mice became moribund, they were euthanized and their brains were harvested for RNA isolation. RABV-N mRNA in the cerebrum, cerebellum, and brain stem was quantified by qPCR. A standard curve was generated from serially diluted plasmids carrying a RABV-N gene and the copy numbers of N mRNA were normalized to 1 mg of total RNA. (* $p < 0.05$, ns means no significant difference).

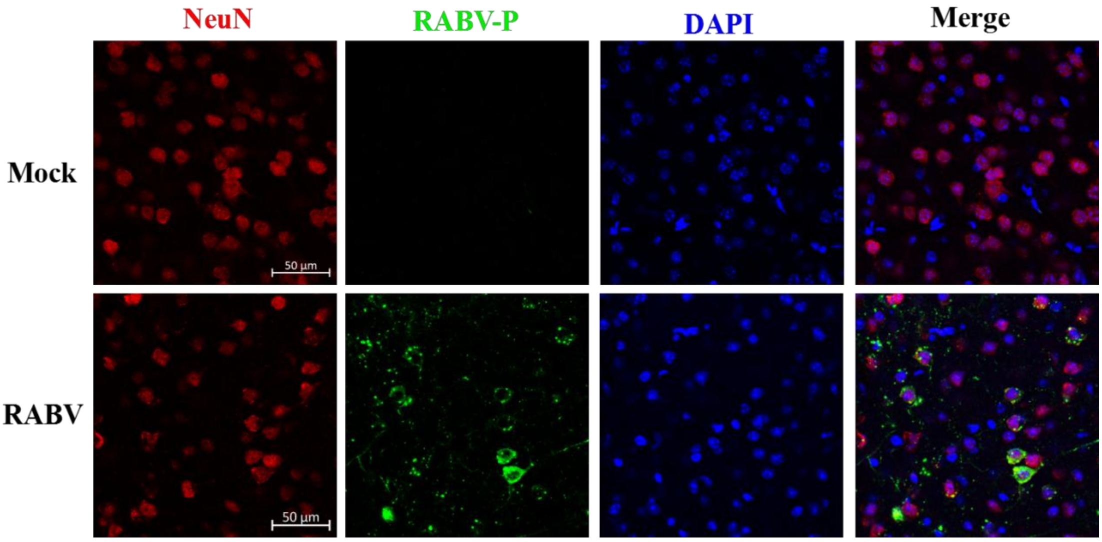


**Additional file 4: Figure S4 IFA staining of RABV-infected neurons in the mouse brain, Related to Figure 3.** C57BL /6 mice (6-8-week-old, n=5) were i.m. inoculated with 10×$LD_{50}$ of RABV-EGFP. When mice became moribund, they were euthanized and their brains were harvested for frozen section preparation. After being stained with antibody against NeuN or RABV-P, the immunofluorescence of frozen sections was observed under an Olympus IX51 microscope. Scale bar, 50 µm.

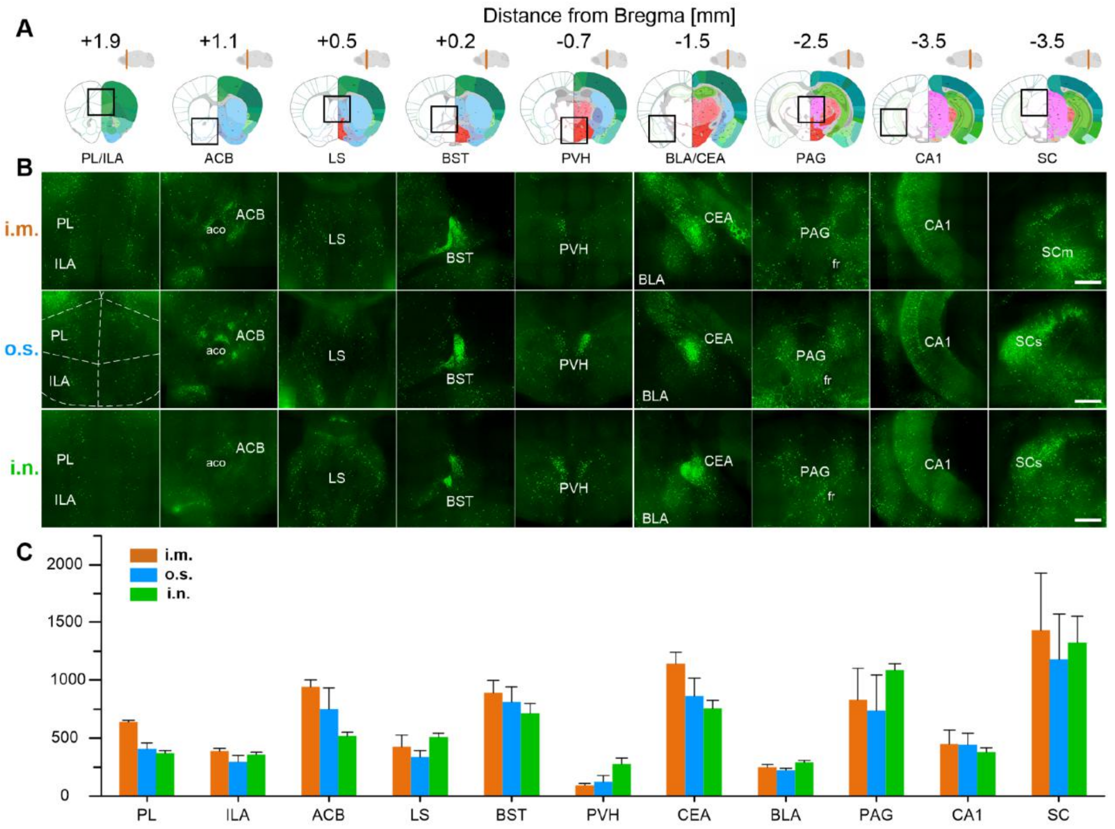


**Additional file 5: Figure S5 Identification of RABV infection in fear-related regions, Related to Figure 3.** (A) The anatomical localization of the selected coronal sections shown in (B) is indicated by black boxes. The distance of the selected coronal section from the bregma is also indicated. (B) Representative pictures of RABV infection in fear-related nuclei. Scale bar, 500 μm. (C) Quantification of RABV-infected neurons in fear-related nuclei (n=3).

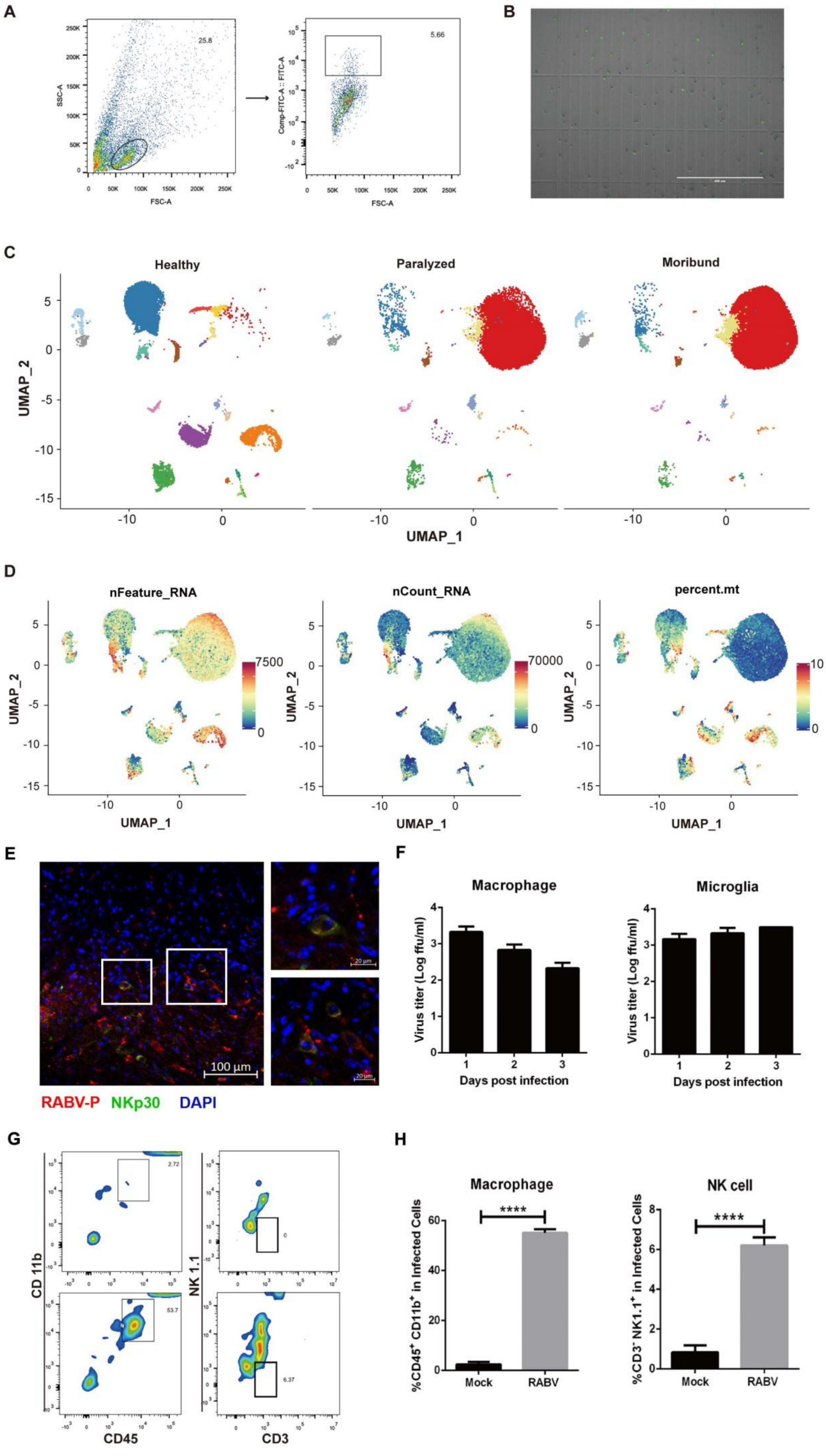
A
SSC-A
FSC-A
25.8
Comp-FITC-A :: FITC-A
5.66
B
C
Healthy
Paralyzed
Moribund
UMAP_2
UMAP_1
D
nFeature_RNA
nCount_RNA
percent.mt
7500
70000
10
E
100 µm
20 µm
RABV-P
NKp30
DAPI
F
Macrophage
Microglia
Virus titer (Log ffu/ml)
Days post infection
G
CD 11b
CD45
NK 1.1
CD3
2.72
53.7
0
6.37
H
Macrophage
NK cell
%CD45+ CD11b+ in Infected Cells
%CD3- NK1.1+ in Infected Cells
****
Mock
RABV

**Additional file 6: Figure S6. Overview clustering results, Related to Figure 4** (A) Gating strategy of flow cytometry. For mice at paralyzed or moribund stage, EGFP-positive cells were enriched by flow cytometry as panel A (right); for healthy mice, single cells were enriched by flow cytometry as panel A (left). (B) Image of EGFP-positive cells sorted by flow cytometry under an immunofluorescence microscope. Scale bar, 400 μm. (C) UMAP projection of three conditions. Each dot corresponds to a single cell, colored according to cell type and the same in Figure 4B. (D) UMAP of UMIs (left), gene counts (middle) and percentage of mitochondrial genes (right) in all cells. (E) RABV-infected NK cell were confirmed by IFA. Mice inoculated with $10\times LD_{50}$ RABV were euthanized with $CO_2$ at the moribund stage, and brains were collected and prepared for IFA staining. The frozen sections were stained with antibodies against RABV-P and NKp20 (NK cell marker), and then the slides were observed under an immunofluorescence microscope. Scale bar, 100 μm. (F) RABV infection in primary microglia and macrophage cultures. Primary microglia and macrophage were prepared and infected with RABV, and then the supernatants were collected for virus titration. (G-H) RABV-infected NK cell and macrophages in the mouse brain were confirmed by flow cytometry. Gating strategy of macrophage (Figure S6G left) and NK cell (Figure S6G right) are shown. C57BL/6 mice (n=5) inoculated with RABV were euthanized at the stage of moribund, and brains were collected and dissociated into single cells. The percentage of macrophage cells (Figure S6H left) and NK cells (Figure S6H right) among RABV-infected cells were calculated by flow cytometry (**** $p<0.01$; Student's *t* test).

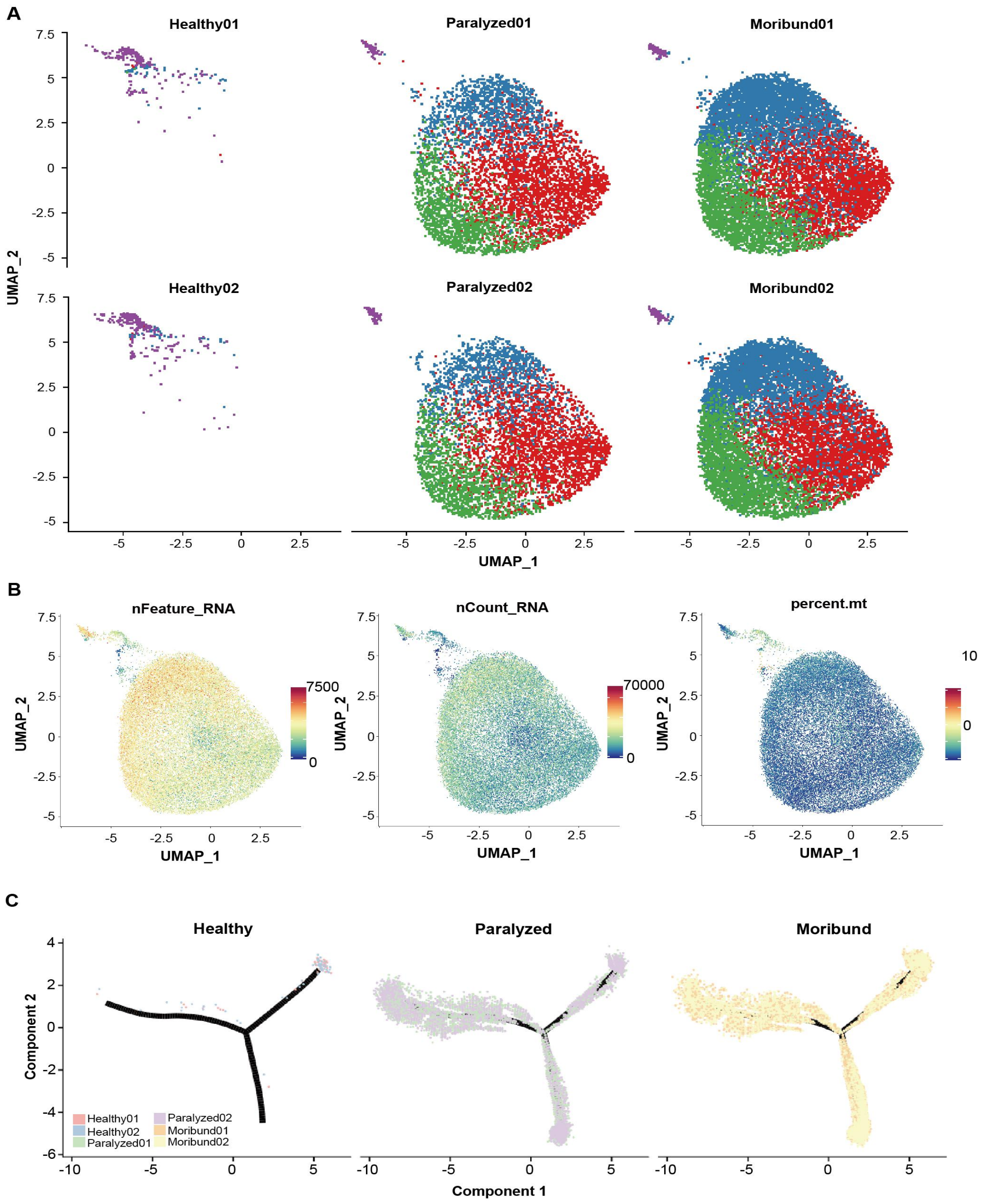


**Additional file 7: Figure S7. Additional features of macrophage subsets, Related to Figure 5 and Figure 6** (A) UMAP projection of each sample. Each dot corresponds to a single cell, colored according to cell type and the same in Figure 5A. (B) UMAP of UMIs (left), gene counts (middle) and percentage of mitochondrial genes (right) in all cells. (C) The potential development trajectory of three macrophage subsets split by conditions. Each dot corresponds to a single cell, colored according to samples.

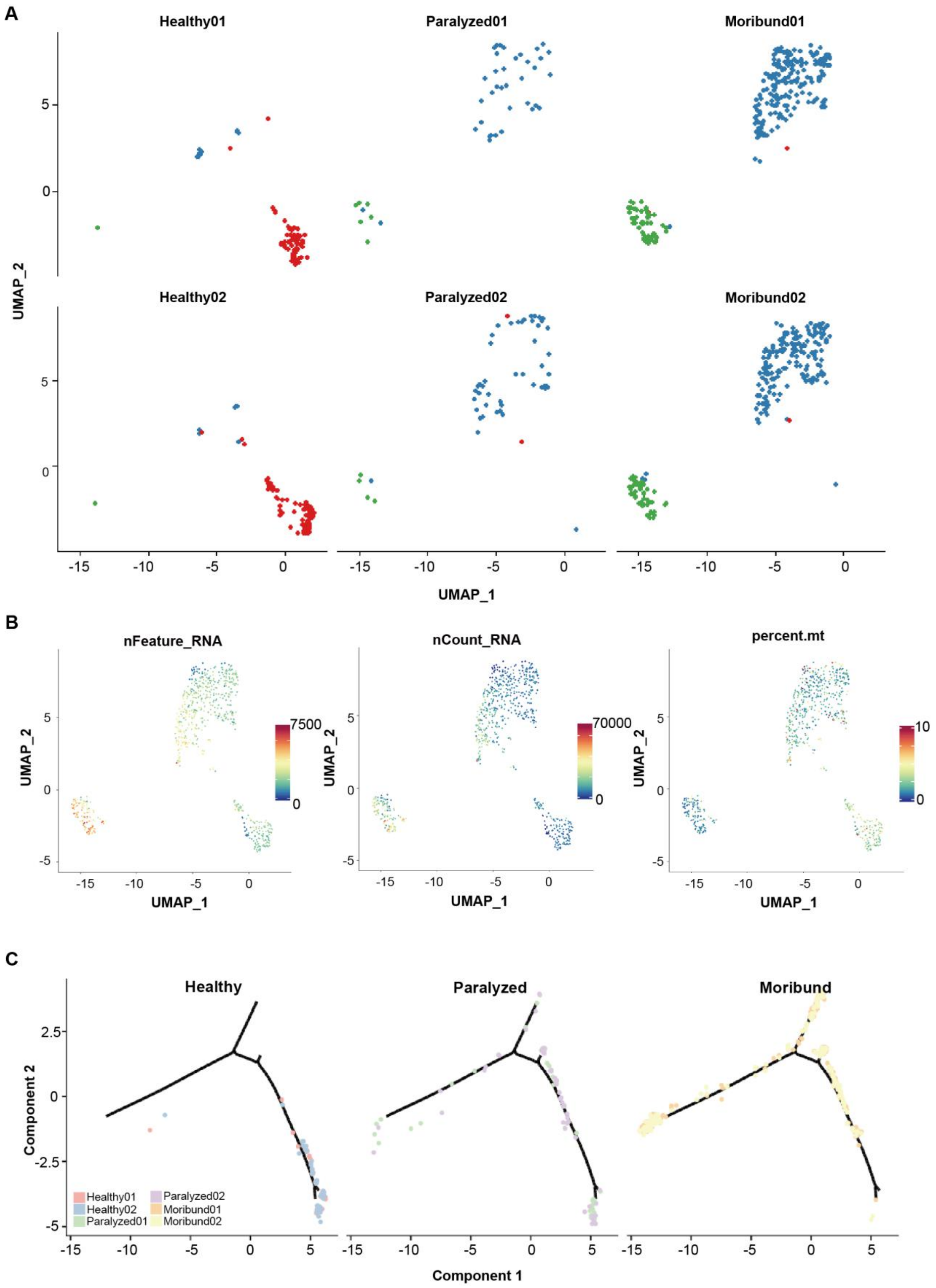


**Additional file 8: Figure S8. Additional features of NK subsets, Related to Figure 7** (A) UMAP projection of each sample. Each dot corresponds to a single cell, colored according to cell type and the same in Figure 7A. (B) UMAP of UMIs (left), gene counts (middle) and percentage of mitochondrial genes (right) in all cells. (C) The potential developmental trajectory of three NK subsets split by conditions. Each dot corresponds to a single cell, colored according to samples.
